# Stretchable, hair-compatible, and long-term stable wearable EEG system

**DOI:** 10.1101/2025.04.28.651043

**Authors:** Ju-Chun Hsieh, Mengmeng Yao, Hussein Mostafa Alawieh, Victoria Koptelova, Satyam Kumar, Kevin Kai Wing Tang, Wenliang Wang, Jinmo Jeong, Hong Ding, Tony Chae, Zoya Ahmad, Derek Wang, Tlali Engrav, Reyna Wang, Ashwin Gupta, Weilong He, William D. Moscoso-Barrera, Ayden Grimes, José del R. Millán, Huiliang Wang

**Affiliations:** Department of Biomedical Engineering, Cockrell School of Engineering, The University of Texas at Austin, Austin, Texas 78712, United States; Chandra Department of Electrical and Computer Engineering, Cockrell School of Engineering, The University of Texas at Austin, Austin, Texas 78712, United States; Department of Neurology, Dell Medical School, The University of Texas at Austin, Austin, Texas 78712, United States; Mulva Clinic for the Neurosciences, The University of Texas at Austin, Austin, Texas 78712, United States

**Keywords:** stretchable, liquid metal, wearable neural interface, electroencephalography, long-term monitoring, hydrogel, brain-computer interface

## Abstract

Electroencephalography (EEG) is a cornerstone in both neuroscience research and clinical diagnostics. However, conventional EEG monitoring faces hardware limitations, particularly its adaptability and stability. Headsets either require complicated wiring or do not have enough stretchability and wearability to comply with the diverse head anthropometry and hair conditions of the user population. Additionally, there is an inherent tradeoff between wet and dry electrodes in capturing high-fidelity signals from hair-covered scalp regions while ensuring continuous and long-term recording quality. Here, we present a Mesh-integrated, Stretchable, and Hair-compatible EEG system engineered to overcome these limitations. By incorporating a kirigami-inspired mesh design and stretchable eutectic Gallium-Indium interconnects, MindStretcH adapts to various head sizes and allows for easy wearing and removal. Moreover, its uniquely designed porous, conical, soft 3D-printed mold, embedded with hydrogel electrodes, effectively penetrates hair layers to deliver low impedance and sustained signal integrity with minimal discomfort. We validate MindStretcH through offline and online EEG-based brain-computer interface tasks over a month, demonstrating its exceptional stability in continuous monitoring and dynamic applications. These results mark a promising advance toward non-invasive neural interfaces in both clinical and everyday use.

## Introduction

Electroencephalography (EEG) has been a cornerstone of neuroscience research and clinical applications for nearly a century.^1,2^ Its outstanding temporal resolution enables real-time monitoring of neural dynamics, rendering it indispensable for the diagnosis and management of neurological disorders, including epilepsy, sleep disorders, and traumatic brain injuries (TBI).^3–5^ In addition, EEG has emerged as a foundational technology in non-invasive brain-computer interfaces (BCIs), facilitating direct communication between the brain and external devices without the necessity of craniotomy. This advancement has broadened its utility beyond traditional medical contexts to encompass applications such as intelligent prosthetic control,^6,7^ communication restoration,^8,9^ and neurorehabilitation training.^10–13^

The gold-standard EEG recording systems typically consist of a heavily wired EEG head cap equipped with either wet or dry electrodes.^14^ However, EEG head caps in commercial space are generally rigid or exhibit limited stretchability. The solutions provided by the existing head caps either incorporate mechanisms to induce stretchability in the headset or offer a range of sizes.^15,16^ This often results in bulky systems or limits the adaptability of the headset to various head shapes and sizes.^17^ These constraints ultimately increase operational costs for facilities engaged in EEG recordings,^18^ particularly when catering to populations with diverse head dimensions, such as infants and adolescents, whose head sizes increase along with their development.^19^ In addition to the adaptability of the head cap, the conundrum of recording at the hair-cover area is also unaddressed. As mentioned, the gold-standard systems typically use either wet or dry Ag/AgCl electrodes.^14,20^ The wet-electrode setting relies on conductive gels to create low-impedance connections with the scalp. Although wet electrodes offer high signal quality, they require preparation by trained personnel and often cause discomfort for users due to the untidy nature of gel application and its propensity to dry out quickly.^21^ On the other hand, the dry counterpart eliminates the need for conductive gels while offering rapid setup and greater mobility.^21^ However, dry electrodes often compromise overall signal quality due to poor electrode-skin contact during movement.^22^ Frequently, dry electrodes require a pointed or conical design to ensure contact with the scalp, particularly in the presence of hair regions.^21^ However, this method often introduces pain and becomes impractical for long-term monitoring.^21^ In addition to the mentioned issues, hair significantly complicates high-fidelity EEG recordings by obstructing the conformability of the head cap and the stable electrode-skin contact. In clinical practice, this issue is commonly mitigated by shaving most of the subject’s hair to ensure reliable recordings—an impractical solution for everyday use that also imposes considerable psychological burdens, particularly on children, adolescents, and female patients.

Recent breakthroughs in skin-conformable electronics have shown promise in addressing this challenge. For example, Tian et al. demonstrated a large-area epidermal electronic interface that is compatible with magnetic resonance imaging (MRI) while providing whole-scalp EEG recordings, enabling multimodal measurements of prosthetic control and cognitive monitoring.^23^ More recently, de Vasconcelos et al. demonstrated a scalp-printed electronic tattoo BCI system, highlighting the potential of digital fabrication techniques for producing ultrathin conformal EEG sensors.^24^ However, while effective, such EEG recordings still require shaving the subject’s hair to facilitate the scalp-printing process. Recent innovations—including novel electrode designs for coarse or curly hair and hydrogel materials with phase-changing properties^25^ that promote the formation of electrodes on the skin—offer promising alternatives.^26,27^ However, it is difficult for electrode formation and removal in the full multi-channel EEG headset in these systems. Developing a one-for-all, convenient-to-wear, high-quality recording and long-term stable multichannel EEG headset remains challenging.

To address these critical gaps present in existing technologies, we have introduced a novel **M**esh-**in**tegrate**d, Stretc**hable, and **H**air-compatible EEG system (MindStretcH) explicitly designed to surmount challenges related to adaptability, comfort, signal fidelity, and long-term usability. **Figures 1a and b** illustrate the users with diverse head sizes wearing MindStretcH. **Figure 1c** illustrates the exploded view of the MindStretcH EEG system. Our system features a Kirigami-inspired mesh design and Gallium-Indium eutectic (EGaIn) stretchable interconnects, resulting in outstanding stretchability and mechanical compliance (**Figure 1d**). This design enables the headset to conform to diverse head sizes while ensuring stable electrode contact quality and signal fidelity. In contrast to conventional designs, this methodology ensures adaptability across diverse conditions and enhances usability in dynamic applications, such as long-term, repeated EEG recordings or EEG-based BCI operations. To mitigate hair-induced signal degradation—a persistent challenge in high-fidelity EEG recordings—We have developed a porous conical soft three-dimensional conical electrode structure integrated with an innovative hydrogel comprising **G**lycerol, potassium chloride (**K**Cl), and 2-Acrylamido-2-methyl-1-propanesulfonic acid sodium salt (**AMP**SNa), collectively referred to as **GKAmp** (**Figure 1e**). This unique design enables the electrodes to penetrate through hair layers (**Figure 1f**) and establish direct contact with the scalp while maintaining low contact pressure for comfortable wearing and low electrical impedance for high-quality recording over extended periods. Unlike traditional hydrogel electrodes that struggle in hair-covered regions or dry environments, our conical soft 3D electrodes provide mechanical compliance that adapts to different scalp morphologies while exerting sufficient contact pressure for stable signal acquisition. Finally, we validate our system through both offline and online longitudinal EEG-based BCI tasks involving neural signal processing and real-time feedback mechanisms. This evaluation not only demonstrates the practicality and longevity of our stretchable EEG headset but also highlights its potential for facilitating advanced EEG-based applications, such as prolonged EEG monitoring and adaptive brain-machine interaction (**Figure 1g**).

**Figure 1.**
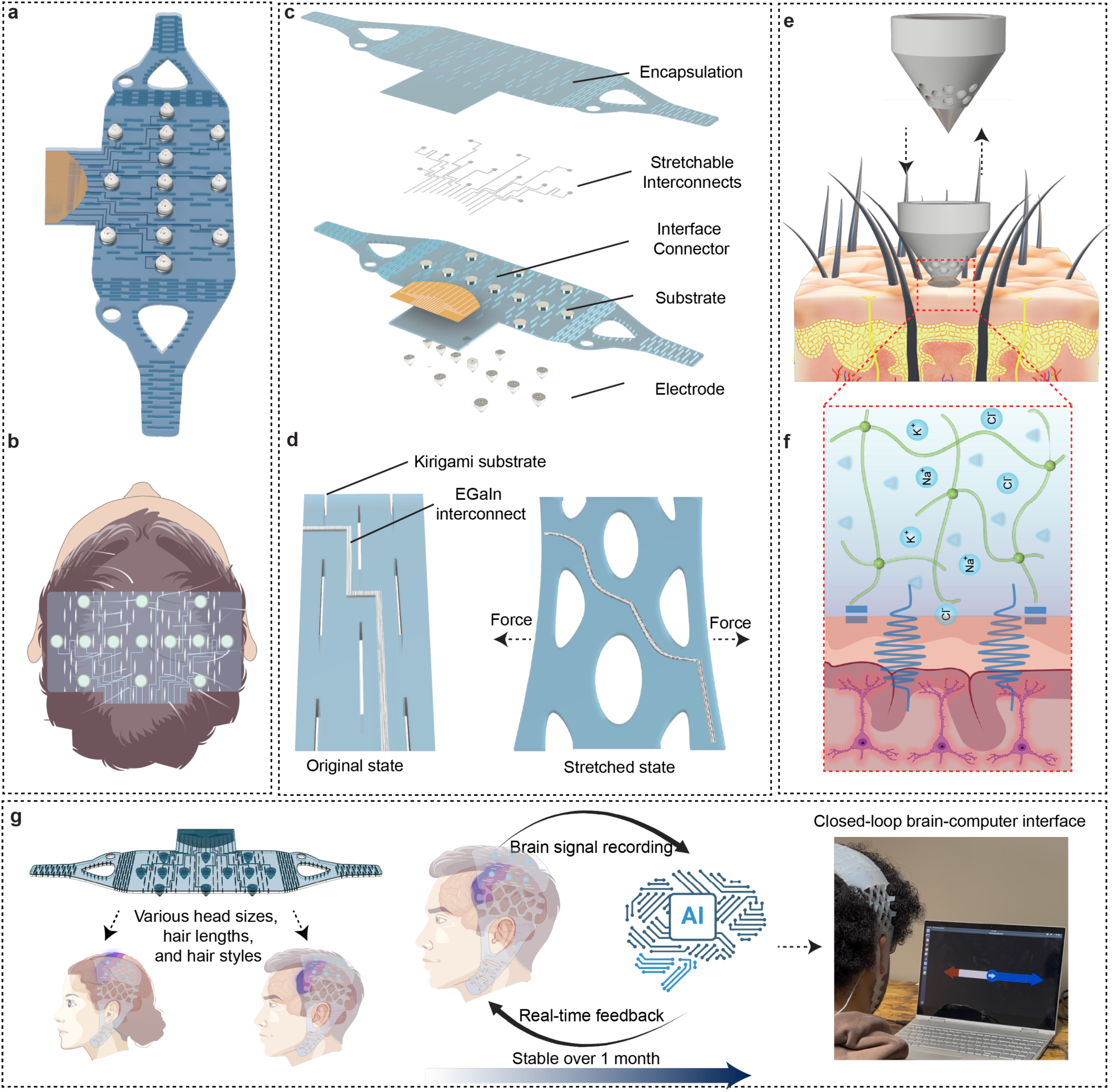
Design and functionality of the mesh-integrated, stretchable, and hair-compatible (MindStretcH) EEG system. **(a)** Top view of the MindStretcH EEG system, showcasing its mesh-integrated design for adaptability to diverse head sizes. **(b)** Illustration of the device positioned on a user’s head. **(c)** Exploded view of the MindStretcH system, highlighting its key components: encapsulation layer, stretchable interconnects, interface connector, substrate, and electrodes. **(d)** Kirigami-inspired mesh structure with EGaIn interconnects in original and stretched states. **(e)** Porous conical soft 3D electrode design integrated with GKAmp hydrogel for effective penetration through hair layers and stable scalp contact. The inset image shows the fabricated electrode. **(f)** Schematic representation of the electrode’s interaction with hair-covered scalp regions. **(g)** Schematic of using the MindStretcH EEG headset for long-term neural signal acquisition to achieve stable closed-loop BCI. The MindStretcH EEG headset is designed to accommodate various head sizes, hair lengths, and hair styles.

## Results

### Design and Performance of MindStretcH

**Figure 2a** presents the top-down view of the MindStretcH EEG system, highlighting its kirigami-inspired mesh design and showing its structural layout and overall electrode placement. The detailed fabrication process is illustrated in **Figure SI1** and described in Methods. MindStretcH’s design integrates a Kirigami-inspired mesh structure with strategically placed cuts to enhance stretchability and mechanical adaptability to control the deformation and localized expansion to conform to varying head sizes. (**Figure 2a and SI2**). Close-up views of the patterned substrate before and after tensile deformation (**Figures 2b and 2c**) demonstrate how the slits expand into a mesh structure under strain, enabling significant mechanical elongation without compromising structural integrity. Kirigami-inspired designs have been widely adopted in stretchable electronics due to their ability to accommodate large deformations while maintaining functional performance.^28–31^ To further illustrate the design rationale, **Figure SI3** provides detailed views of two Kirigami patterns used in different regions (the temporal region in **Figure SI3a**, and the scalp region in **Figure SI3b**) in MindStretcH. The pattern that sits at the temporal region (**Figure SI3a**) features parameters of horizontal spacing (L_HS_) = 5 mm, vertical spacing (L_VS_) = 2.5 mm, and cut length (L_CL_) = 20 mm, resulting in an aspect ratio of 4.0. This higher aspect ratio ensures greater stretchability to conform to complex contours around the area of the head while maintaining comfort. In contrast, the pattern that sits at the scalp region (**Figure SI3b**) uses parameters of L_HS_ = 10 mm, L_VS_ = 5 mm, L_CL_ = 15 mm, yielding an aspect ratio of 1.5. This lower aspect ratio ensures that the headset remains securely fitted while accommodating less deformation compared to the former one.

**Figure 2:**
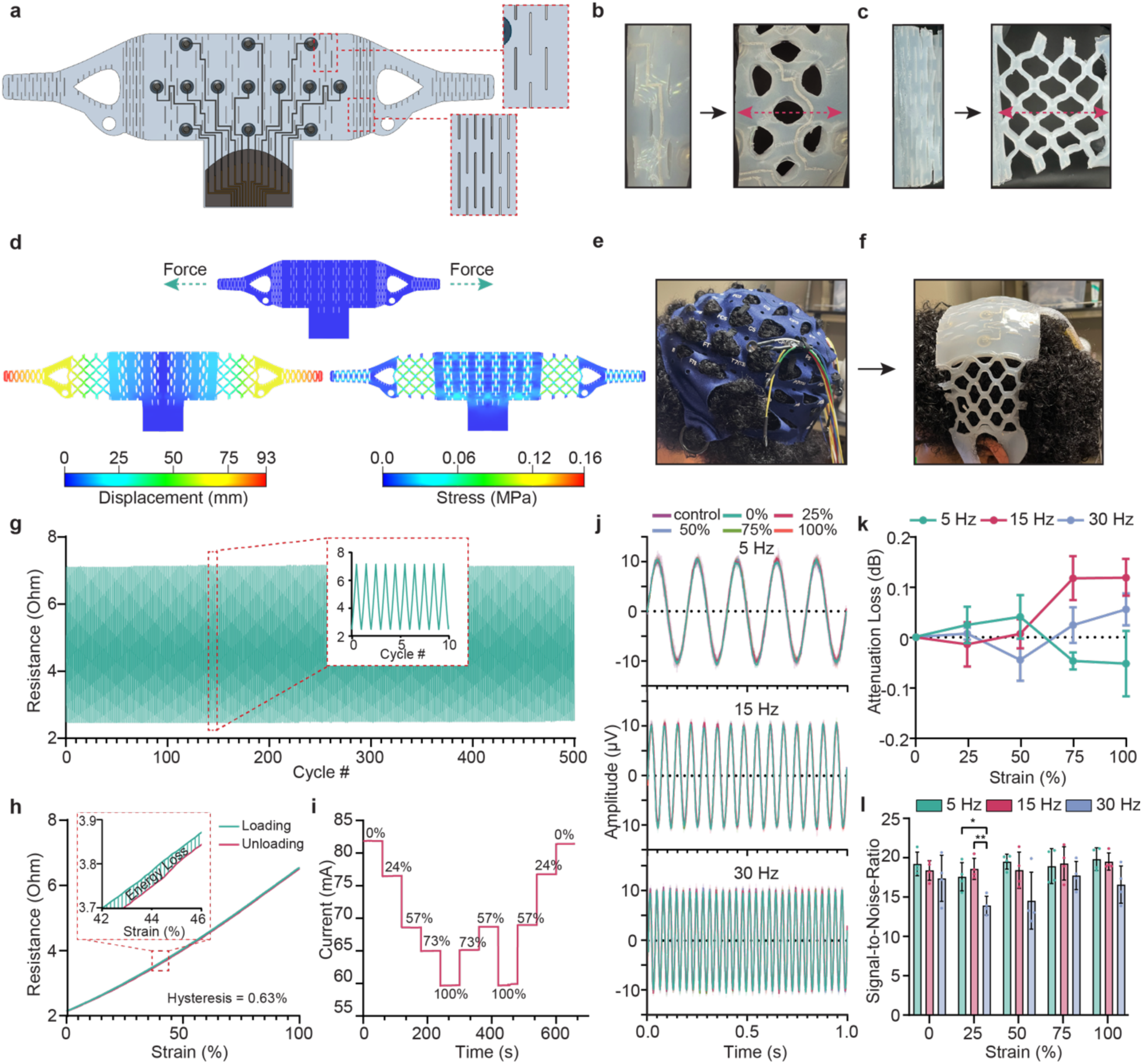
Design, mechanical characterization, and electrical performance of the MindStretcH wearable EEG system. **(a)** Top-down view of the MindStretcH headset, showing the electrode layout and interconnect design. The zoomed-in sections highlight the kirigami-inspired patterns integrated into the design to enhance mechanical stretchability. **(b, c)** Close-up views of the kirigami-inspired patterns before and after stretching. **(d)** Finite element analysis (FEA) simulation results of the MindStretcH structure when stretched to fit an average ear-to-ear distance of 40 cm. Top: the original state. Bottom Left: Displacement distribution (mm) in the stretched state. Bottom Right: Stress distribution (MPa) in the stretched state. **(e, f)** Comparison of conventional EEG caps **(e)** and the MindStretcH device **(f)** when fitted to a subject. The conventional cap fails to conform to head morphology, whereas MindStretcH adapts seamlessly. **(g)** Resistance of EGaIn interconnects during 500 loading and unloading cycles at 100% strain. Inset: Magnified view of several cycles showing consistent resistance behavior. **(h)** The resistance-strain relationship of EGaIn interconnects during a single loading-unloading cycle, generating a minimal hysteresis of 0.63%. **(i)** Current flow through the EGaIn interconnects over time under various strain levels (0%, 24%, 57%, 73%, and 100%), demonstrating stable electrical performance under stretching. **(j)** Signal fidelity of EGaIn interconnects under strain conditions (0%, 25%, 50%, 75%, and 100%), compared to control signals from traditional metal wires, across three representative frequencies (5 Hz, 15 Hz, 30 Hz). Overlaid waveforms exhibit minimal distortion. **(k)** Attenuation loss across three frequencies (5 Hz, 15 Hz, and 30 Hz) as a function of strain. Attenuation remains below 0.2 dB, ensuring reliable signal transmission. **(l)** Signal-to-noise ratio (SNR) across strain levels for three frequencies (5 Hz, 15 Hz, and 30 Hz). The EGaIn interconnects consistently generated an SNR of ∼13 or above with minor deviations.

To evaluate the mechanical performance of MindStretcH, we conducted finite element analysis (FEA) simulations under realistic stretching conditions. **Figure 2d** illustrates the simulation results of displacement (bottom left) and stress (bottom right) distributions, where the headset is stretched under a simulated tensile loading scenario to generate controlled expansion to fit an ear-to-ear distance of 40 cm. The results confirm that each EEG channel repositions effectively to approximate the selected electrode locations in the international 10-10 system, ensuring proper neurophysiological mapping during use. The corresponding stress distribution (**Figure 2d, bottom right**) reveals stress concentration primarily near the edges of the slits. However, maximum stress values remain well below Ecoflex-0030’s tensile strength (∼1.38 MPa), ensuring structural integrity under applied loads. The Kirigami patterns delocalize stress across multiple deformation points. The higher L_VS_ increases critical buckling force, while longer L_CL_ enhances extensibility. The design for the temporal region’s higher aspect ratio (L_CL_/L_HS_=4.0) facilitates greater elongation compared to the scalp region (L_CL_/L_HS_=1.5), which prioritizes stability. These FEA results align well with the findings in a prior effort by Shyu et al., 2015, which demonstrated that kirigami patterns significantly improve strain capabilities in materials by delocalizing stress over numerous preset deformation points^29^. **Figures 2e-f** compare MindStretcH with a conventional EEG cap on a subject with challenging head morphology and hair conditions. The conventional cap fails to conform effectively due to limited stretchability, resulting in poor electrode contact (**Figure 2e)**. In contrast, MindStretcH demonstrates superior adaptability, maintaining secure coverage regardless of head size or hair type (**Figure 2f**). This adaptability addresses a key limitation of traditional EEG caps.^14^

To characterize the electrical performance of the Gallium–Indium eutectic (EGaIn) liquid metal interconnects integrated within the MindStretcH device, we conducted a series of tests to evaluate signal consistency and fidelity under varying deformation conditions (stretching, twisting, and bending).^32^ To start, we characterize the electromechanical properties of EGaIn interconnects. The experimental setup and circuit diagram are shown in **Figures SI4 and SI5**, respectively. **Figure 2g** shows the resistance stability during 500 cycles of loading and unloading at 100% strain, with the inset magnifying the view of ten cycles to show the detailed signal quality. The results reveal consistent resistance fluctuations within a narrow range, underscoring excellent cyclic durability. Similarly, **Figure 2h** depicts the linear resistance-strain relationship during a complete loading-unloading cycle, with a minimal hysteresis of 0.63%, indicating reliable electromechanical coupling and elastic recovery. Furthermore, **Figure 2i** evaluates the current response under randomized strain levels (0%, 24%, 57%, 73%, and 100%), confirming stable current flow with minimal fluctuations. These results collectively establish the mechanical resilience of EGaIn interconnects, ensuring consistent electrical performance under dynamic strain conditions, a critical feature for wearable EEG systems. We further assessed the signal fidelity of MindStretcH under mechanical deformation, including varying strain conditions (**Figures 2j-l**). The picture of the experimental setup and the circuit diagram are shown in **Figures SI6 and SI7**, respectively. The image of the experimental setup to achieve all deformation conditions is shown in **Figure SI8**. The control signal of interest was set to have a peak-to-peak amplitude of 20 µV to mimic the typical amplitude of EEG signals. **Figure 2j** compares the signal fidelity of three different signal conditions (from top to bottom, 5 Hz, 15 Hz, 30 Hz) to control metal wires at various strain percentages (0%, 25%, 50%, 75%, and 100%). The overlaid waveforms exhibit consistent signal amplitude and phase alignment, confirming their ability to transmit signals with high fidelity across deformation states. **Figure 2k** quantifies attenuation loss for three representative EEG frequencies (5 Hz, 15 Hz, and 30 Hz), showing minimal attenuation (<0.2 dB) under all strain conditions. This ensures reliable signal acquisition of weak electrophysiological signals such as EEGs. Lastly, **Figure 2l** presents the signal-to-noise ratio (SNR) under the same conditions, with values consistently exceeding 13, and shows no statistically significant difference across strain conditions, except for a 30 Hz signal at 25% strain. This phenomenon can be attributed to the intrinsic characteristics of signal transmission systems, wherein higher frequencies exhibit increased susceptibility to attenuation, due to resistive and capacitive effects present within the transmission medium.^33^ This demonstrates the interconnects’ robustness in preserving signal clarity even at 100% strain. In addition to strain tests, we evaluated the performance of the EGaIn interconnects under twisting conditions from 0 to 5 twists (**Figure SI9**) to evaluate their suitability for wearable EEG systems in dynamic real-world applications. As shown in **Figure SI9a**, the overlaid waveforms for 5 Hz, 15 Hz, and 30 Hz remain consistent in amplitude and phase alignment across most of the twist counts, indicating stable signal transmission under rotational deformation. The attenuation loss is below 0.2 dB, emphasizing the robust signal transmission (**Figure SI9b**). The interconnects maintain a consistent SNR above 15 across increasing twist counts, with statistically significant differences observed only in a few conditions (**Figure SI9c**). Similarly, we examined the bending deformation at angles ranging from 0° to 160° (**Figure SI10**). As observed in **Figure SI10b**, attenuation remains minimal (<0.2 dB), even at the most extreme bending angles. Finally, **Figure SI10c** indicates that the SNR remains above 15 across most bending angles, with statistically significant differences observed only at 45° and 160° for specific frequencies (5 Hz and 30 Hz, respectively). These results suggest that the EGaIn interconnects and MindStretcH are capable of transmitting electrophysiological signals without significant loss of fidelity, even under conditions of severe mechanical deformations.

### Integration of Hydrogel and 3D-Printed Elastic Electrodes for Hair-Compatible EEG Sensing

We developed an integration of hydrogel and 3D-printed conical elastic electrodes for the MindStretcH system, addressing hair interference and prolonged wear stability challenges. Our previous work developed an injectable and spontaneously cross-linked AIRTrode hydrogel that exhibits low impedance and high signal fidelity over extended periods.^27^ However, the dark and opaque color of the AIRTrode hydrogel may raise potential FDA regulation concerns. In addition, the presence of PEDOT:PSS may limit the overall shelf lifetime of the hydrogel and the integrated devices.^34^ Here, based on the foundation of AIRTrode, a novel iteration of ionic hydrogel, GKAmp, is presented (**Figure 3a**). GKAmp hydrogel comprises **G**lycerol, potassium chloride (**K**Cl), and 2-Acrylamido-2-methyl-1-propanesulfonic acid sodium salt (**AMP**SNa). GKAmp’s highly dissociative sulfonate groups form a dense network of mobile charge carriers, ensuring stable ionic conduction. Moreover, the hydroxyl and sulfonate groups establish abundant hydrogen bonds with skin amines, contributing to strong adhesiveness, which is particularly beneficial for maintaining stable contact in regions with dense hair coverage.^22,27,35^ Glycerol forms a binary solution with the hydrogel’s water content,^27^ minimizing evaporation and thereby prolonging both the electrical and mechanical stability of GKAmp over time. Furthermore, the KCl regulates ionic strength, maintaining a consistent Debye screening length to optimize charge transfer dynamics and prevent interface polarization.^36,37^ As shown in **Figures 3b**, **SI11, and SI12**, GKAmp adheres readily to various skin surfaces, including the fingers and forearm, with a maximum adhesion strength of 1.56 N/cm. Notably, even after eight repeated adhesion cycles, the hydrogel maintains its stability, demonstrating excellent reusability. Moreover, GKAmp features tunable mechanical properties, with a tensile strength of 32.95 kPa, an elongation at break of 877%, and an average Young’s modulus of 16.5 kPa, ensuring excellent compliance and conformability at the skin interface (**Figure 3c and SI13**). When compressed to 30% of its original thickness, the hydrogel’s compressive stress increases from 0.23 N ± 0.10 kPa to 1.19 N ± 0.19 kPa, indicating its ability to accommodate various scalp conditions through controlled crosslinking adjustments (**Figure SI14**). We analyzed the impedance-frequency response of different electrode types to assess their suitability for bioelectrical signal acquisition (**Figure SI15**). GKAmp and semi-dry electrodes (marked as “Semi-dry1” and “Semi-dry2” in **Figure 3d**) showed lower impedance compared to dry electrodes but slightly higher than wet electrodes, with stable performance across frequencies. The impedance decreased with increasing frequency, consistent with capacitive effects (**Figure 3d**). We observed that the interfacial impedance of GKAmp hydrogel decreased significantly with increasing KCl concentration (0.29 wt%, 0.58 wt%, and 0.88 wt%) (**Figures 3e and SI16**). This trend aligns with the ion compensation effect of KCl in the hydrogel matrix, where increasing KCl concentration provides more mobile ions to facilitate charge transfer, particularly in the low-frequency range (<10 Hz), where ionic conduction dominates. More importantly, the hydrogel maintained minimal degradation in charge transport properties over four weeks (**Figure 3f and SI17**), suggesting its suitability for prolonged EEG monitoring. The weight retention analysis over 600 hours further revealed that glycerol effectively modulated water loss kinetics, maintaining hydration levels above 80% compared with hydrogels without glycerol, despite fluctuations in environmental humidity (30–55%) (**Figure 3g and SI18**). We conducted cyclic charge injection capacity (CIC) testing over 1500 cycles. As shown in **Figure 3h**, the GKAmp hydrogel maintained a stable charge injection capacity over four weeks, indicating sustained electrochemical performance (**Figure 3i**). This stability is attributed to the optimized ionic conductivity facilitated by KCl incorporation and the hydrogel’s hydration retention properties, which together ensure efficient charge transport at the skin-electrode interface. Furthermore, cell cytotoxicity tests showed that GKAmp hydrogel electrodes maintained high cell viability (>75%) across all tested KCl concentrations (0.29 wt%, 0.58 wt%, and 0.88 wt%) after 24h and 48h incubation, confirming good biocompatibility (**Figure 3j** and **SI19**). We conducted eyes-open and eyes-closed tests to compare the electrophysiological signal accuracy of GKAmp hydrogel and commercial EEG gel. As shown in **Figure SI20**, both materials demonstrated comparable mu band amplitude (**Figure SI20a**) and power (**Figure SI20b**), with high correlation coefficients for amplitude (r=0.8062) and power (r=0.9408). Time-frequency spectrograms (**Figure SI20c**) showed consistent mu-band patterns between eyes-open and eyes-closed states. These findings confirm that GKAmp hydrogel provides equivalent signal fidelity as commercial EEG gels. To address the limitations of commercial wet EEG electrodes, particularly their restricted applicability to diverse hair conditions and the excessive residual gel left on hair after use, we developed a flexible, 3D-printed conical support platform for GKAmp hydrogel. By functioning as an insulating layer for the GKAmp hydrogel, this platform design effectively prevents the substantial surface area of the internal load from adhering to strands of hair, such that only the hydrogel tip remains exposed, thus creating an electrode suitable for use with hair-compatible hydrogel materials. By utilizing the excellent cohesion of GKAmp and the design of this platform, this electrode can navigate through the hair and establish direct contact with the scalp, regardless of hair density or texture (**Figure 3k and SI21**). To enhance wearing comfort, we introduced a porous structure at the tip of the conical mold. We hypothesized that this porous design would significantly reduce localized stress on the scalp, thereby improving long-term comfort during extended EEG recordings (**Figures 3k-3n**). Both Finite element simulations and experimental results confirmed this hypothesis. Under compression, the porous electrode induced substantially lower surface stress on the scalp compared to its non-porous counterpart under identical external force conditions. The porous structure allows for greater deformation of the electrode, increasing the effective contact area and distributing stress more evenly across the scalp surface (**Figure 3l**) Additionally, displacement simulations **(Figure 3m**) revealed that the porous electrode caused less displacement on the scalp than the non-porous electrode, particularly at the contact periphery. This indicates better stress distribution and reduced pressure points. Experimental compression tests at 10% and 30% strain further validated these findings (**Figure 3n**). At both strain levels, the porous platform exhibited significantly lower stress compared to the non-porous platform, effectively reducing localized pressure on the scalp. Two-way repeated measures ANOVA corroborated the significant differences between porous (10% strain: 1.4 N, 30% strain: 4.0 N) and non-porous (10% strain: 2.2N, 30% strain: 6.3 N) platforms across both strain levels (10% strain: p = 0.0047, n = 4; 30% strain: p < 0.0001, n = 4). As **Figure SI22** shows, the porous and non-porous platforms were tested on a human volunteer’s arm (**Figures SI22a and SI22b**) and forehead (**Figures SI22c and SI22d**) to evaluate residual redness after 10 minutes of wear. The non-porous platform consistently left more pronounced redness on the skin compared to the porous platform, indicating higher localized pressure. In contrast, the porous platform exhibited reduced redness, suggesting that it distributed pressure more evenly across the contact area. This observation aligns with the finite element simulation and experimental compression test results, further validating the hypothesis that the porous design minimizes localized stress and enhances comfort during prolonged use. By combining this porous, 3D-printed elastic conical platform with GKAmp hydrogel, we successfully addressed key shortcomings of commercial wet electrodes. This innovative design ensures stable skin-electrode contact and enhances wearing comfort over extended periods while preventing gel residue on hair. These advancements allow our EEG system to be highly suitable for diverse hair conditions and prolonged EEG recordings.

**Figure 3.**
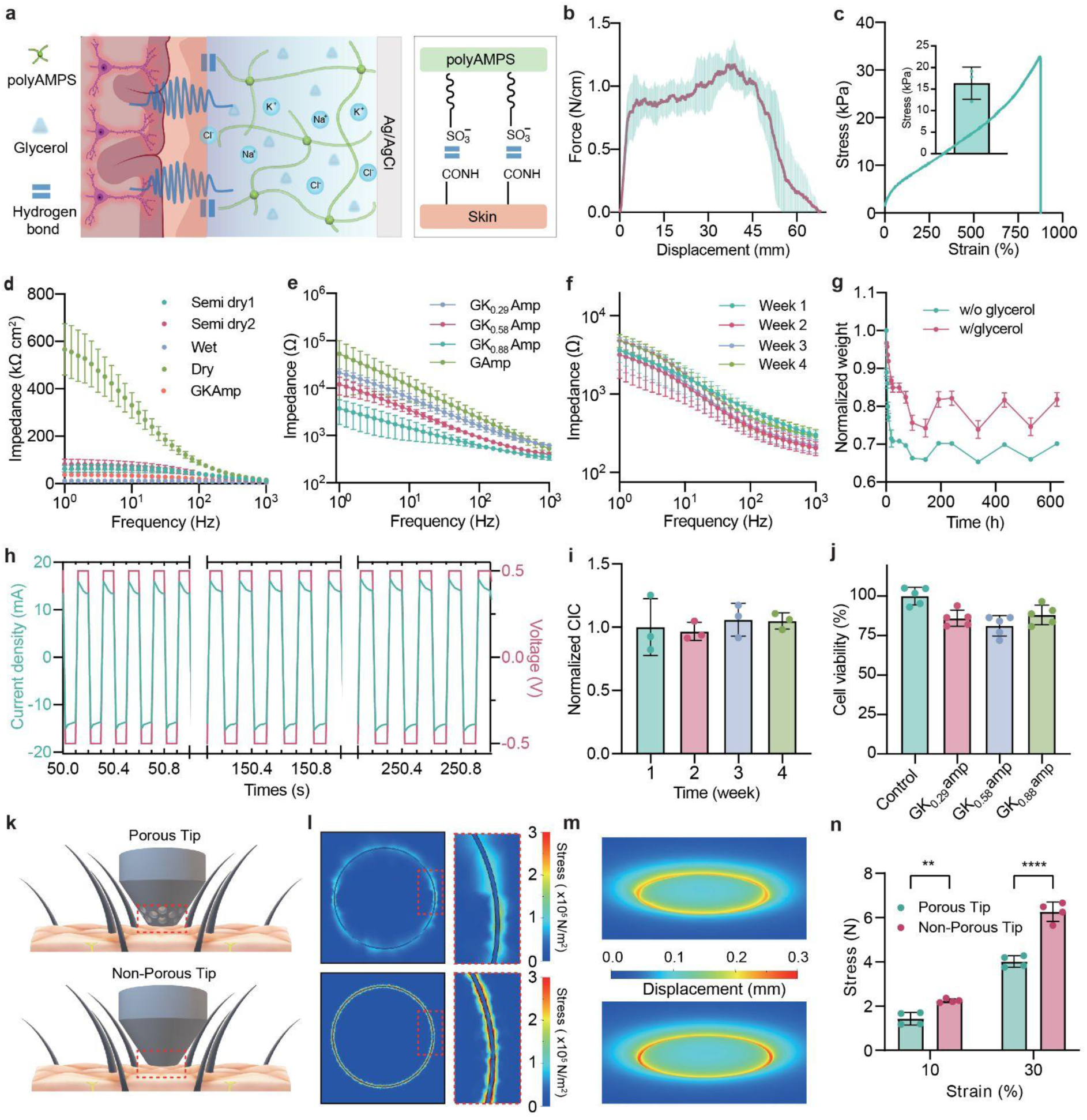
Characterization of the GKAmp hair-compatible electrode for wearable EEG applications. **(a)** Schematic diagram of the hydrogel network in contact with the scalp and electrodes. **(b)** 90-degree peel test results of hydrogel on the skin surface (n=8). **(c)** Stress-strain curves and average Young’s modulus results for hydrogels. **(d)** Comparison of skin impedance between GKAmp and commercial dry and wet electrodes. Impedance spectra of GKAmp hydrogels **(e)** with 0.29 wt% (GK_0.29_Amp), 0.58 wt% (GK_0.58_Amp), and 0.88 wt% (GK_0.88_Amp) KCl and **(f)** over four weeks of measurements. **(g)** Normalized weight retention of hydrogels with and without glycerol over 600 hours. **(h)** Cyclic charge injection capacity (CIC) test GKAmp hydrogels over 1500 cycles **(i)** Normalized CIC of GKAmp hydrogels over four weeks. **(j)** Cell viability analysis for GKAmp hydrogels with different KCl concentrations after 24 h culture of NIH/3T3 fibroblasts. **(k)** Schematic illustration of the porous (top) and non-porous (bottom) conical electrode platforms interacting with the scalp through the hair. **(l)** Finite element simulation results showing stress distribution on the scalp for porous (top) and non-porous (bottom) electrodes under compression. The porous electrode exhibits a broader, more evenly distributed stress pattern compared to the concentrated stress of the non-porous electrode. **(m)** Finite element simulation of displacement on the scalp under compression for porous (top) and non-porous (bottom) electrodes. The porous electrode induces less displacement at the contact periphery, demonstrating improved stress distribution. **(n)** Experimental results comparing stress levels induced by porous and non-porous platforms at 10% and 30% strain. The porous platform shows significantly lower stress at both strain levels (p = 0.0047 at 10% strain; p < 0.0001 at 30% strain), supporting its ability to reduce localized pressure on the scalp for enhanced comfort during prolonged use. Error bars represent mean ± SD; statistical significance is indicated as **p < 0.01, ****p < 0.0001.

### MindStretcH Enables Long-Term and Reliable Motor Imagery EEG Monitoring

To corroborate the feasibility and robustness of the MindStretcH device to support EEG-based brain-computer interface over an extended period, we conducted one-month-long motor-imagery EEG recordings on 6 subjects. Motor-imagery (MI) is a well-established BCI paradigm that elicits reproducible event-related desynchronization (ERD) in sensorimotor rhythms (mu and beta bands), enabling non-invasive detection of cortical engagement during mental rehearsal of movements.^38,39^ This approach is particularly relevant for longitudinal neurorehabilitation applications, where sustained MI practice can drive neural plasticity and functional recovery in motor-impaired populations.^12,39^ **Figure 4a** illustrates a subject wearing the MindStretcH EEG device to perform a motor-imagery task in the experiment. **Figures 4b and 4c** show the grand average normalized power of the mu-band (8–13 Hz) and beta-band (15–30 Hz) over time during the MI task. The shaded regions represent the standard error of the mean (SEM), indicating the consistency of trends across subjects. For both frequency bands, a clear ERD is observed during the MI task, characterized by a power decrease that is sustained throughout the task period. Interestingly, in the first two sessions (first week), the ERDs in the mu band are relatively weaker than in the last three sessions. This can be attributed to the gradual improvement of subjects’ motor-imagery technique.^40^ Overall, these dynamics suggest consistent engagement of the sensorimotor cortex during MI tasks. **Figures 4d–f** examine EEG activity during a resting condition, where participants were instructed to remain relaxed without engaging in motor imagery. **Figure 4d** illustrates a subject wearing the MindStretcH device while performing the rest task. **Figures 4e and 4f** present the grand average normalized power of the mu-band and beta-band, respectively, during rest. Unlike during MI tasks, no ERD is evident in either frequency band during rest, as expected. The stability of mu- and beta-band power across time highlights that resting-state EEG signals remain consistent and unaffected by external tasks, further validating the reliability of the MindStretcH device for capturing baseline neural activity.

**Figure 4:**
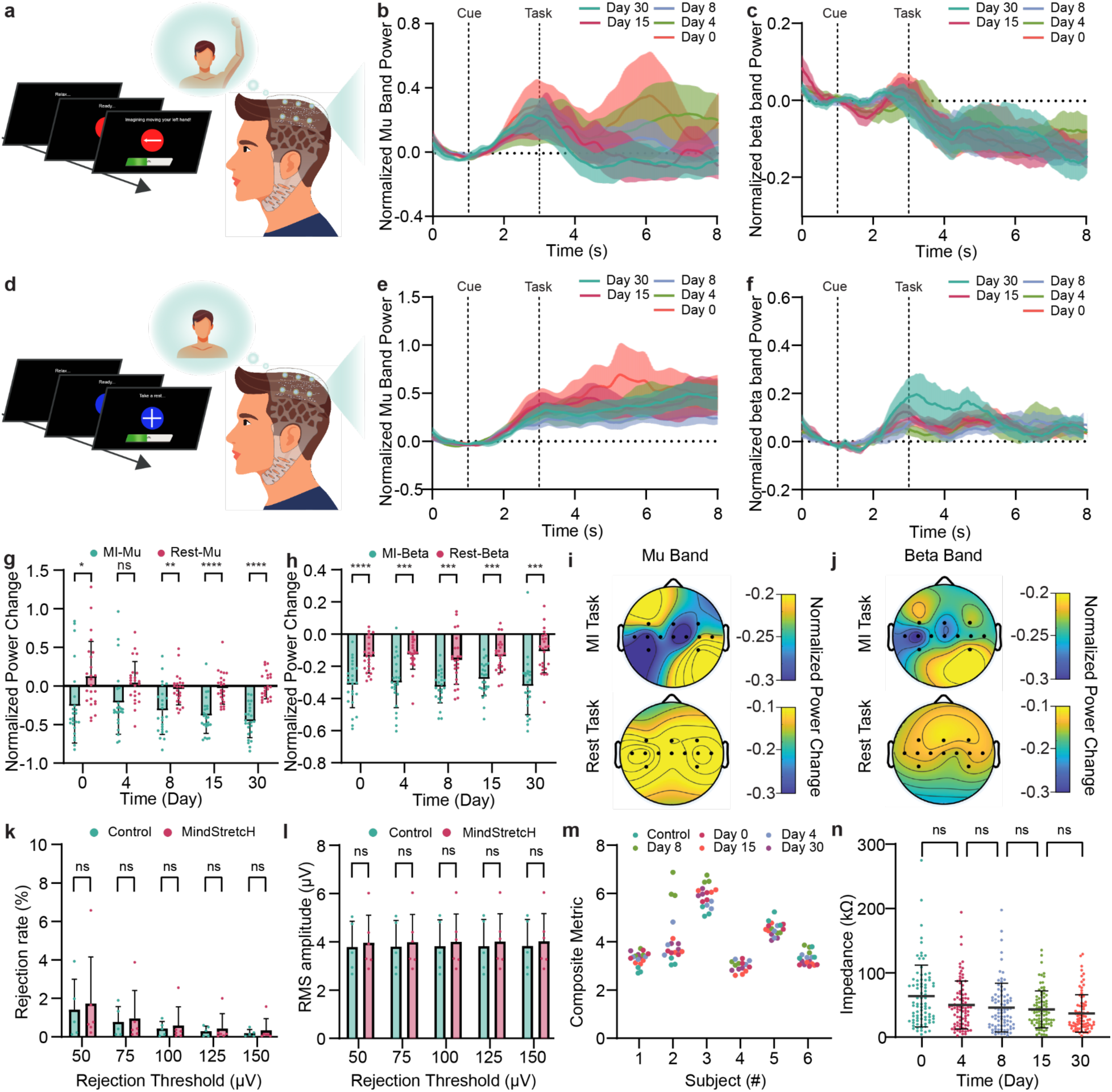
Evaluation of motor imagery (MI) EEG recording and signal quality using the MindStretcH device over a month. **(a)** Schematic illustration of a subject wearing the MindStretcH device during an offline binary-class motor imagery task (Tasks: dominant-hand movement imagery or rest). **(b)** Grand average normalized power of the mu-band (8–13 Hz) across subjects over time (0–8 s) during the last second of the fixation period, cue (2 seconds), task (shown 5 seconds), with shaded regions representing standard error of the mean (SEM). **(c)** Grand average normalized power of the beta-band (18–30 Hz) under identical conditions as **(b)**, with SEM denoted as shaded regions. **(d)** Schematic illustration of a subject wearing the MindStretcH device during an offline rest task, where no motor imagery is performed, and the subject remains in a relaxed state. **(e)** Grand average normalized power of the mu-band (8–13 Hz) across subjects over time (0–8 s) during the last second of the fixation period, cue (2 seconds), and rest task (shown 5 seconds), with shaded regions representing SEM. **(f)** Grand average normalized power of the beta-band (18–30 Hz) under identical conditions as **(e)**, with SEM denoted as shaded regions. **(g)** Normalized Event-Related Desynchronization (ERD) in the mu-band over time at electrode C3/C4 (depending on dominant hand) during motor imagery (MI-mu) and rest (Rest-mu), averaged across 6 subjects, 4 runs per session, and 5 sessions over 30 days. Statistical significance across sessions is marked (ns: not significant; *p < 0.05; **p < 0.01; ***p < 0.001, ****p < 0.0001). **(h)** Normalized ERD in the beta-band under identical conditions as **(d)**. **(i)** Grand-averaged (across all subjects and sessions) topographical map of mu-band ERD during MI tasks (top) and rest (bottom). **(j)** Topographical map of beta-band ERD under similar conditions as **(i)**. **(k)** Subject-wise trial rejection rates at varying thresholds (50–150 µV), comparing the MindStretcH device and a gold-standard EEG cap. Statistical analysis revealed no significant differences between devices across thresholds. **(l)** Root-mean-square (RMS) amplitude of clean trials at varying thresholds, comparing MindStretcH and the gold-standard cap across 6 and 5 subjects, respectively. No significant differences were observed between devices. **(m)** Composite metric scores for signal quality across five sessions for both systems; lower values indicate better signal quality, with no significant differences observed between systems or sessions. **(n)** Longitudinal scalp-electrode impedance (kΩ) for MindStretcH over 30 days. Error bars represent the mean, with no significant changes over time.

The normalized ERD for the mu- and beta-bands at the C3/C4 electrode sites (dependent on the subject’s dominant hand) across the one-month experiment are shown in **Figures 4g and 4h**. The temporal progression of mu-band ERD during motor imagery (MI) tasks reveals significant changes over the one-month experiment, underscoring the specificity and reliability of task-related cortical modulation captured by the MindStretcH device. Statistical analysis using two-way repeated measures ANOVA revealed significant effects for the interaction between conditions (MI vs. Rest) and session progression (p=0.0377), as well as for condition (MI vs. Rest, p=0.0276) and session progression (p<0.0001). These findings highlight both the immediate engagement of sensorimotor cortical areas during motor imagery and the progressive enhancement of cortical modulation over time. Post hoc (Turkey’s test) comparisons demonstrated significant differences in ERD between MI-mu and Rest-mu across most sessions. Notably, on Day 0 (p=0.0258), naive users exhibited a clear differentiation between MI and rest conditions, indicating their ability to engage sensorimotor areas from the outset. However, by Day 4 (p=0.0564, ns), this differentiation became less pronounced, likely reflecting early variability in task performance. From Day 8 (p=0.0043) onward, the significance and magnitude of mu-band ERD steadily increased, reaching peak differentiation by Day 30 (p<0.0001). Interestingly, in the first two sessions, the duration of mu-band ERD was relatively shorter and exhibited a bounded recovery in power during the task phase, as illustrated in **Figure 4b**. This phenomenon is likely attributable to several factors. Similar to motor learning, early sessions may have imposed higher cognitive demands on naive users, leading to inconsistent engagement of sensorimotor areas and shorter ERD durations. As the experiment progressed, neuroplastic adaptation likely facilitated the development of task-specific cortical pathways, enabling sustained ERD during motor imagery. Furthermore, improvements in task familiarity and the refinement of mental imagery strategies contributed to the observed trends.^41^ These changes are corroborated by the topographical maps in the Supporting Information (**Figures SI23 and SI24** for mu- and beta-band, respectively), which illustrate a shift from broader, less distinct sensorimotor activation in early sessions to more localized contralateral activation in later sessions. This spatial refinement reflects the progressive development of task-specific neural efficiency over time. These results hint to underlying cortical plasticity mechanisms involved in learning the control of BCIs with longitudinal training, a well-documented phenomenon in MI-based BCI studies^42–45^. The beta-band ERD also exhibited task-related significant difference, with effect of conditions (motor imagery vs. rest, p=0.0276) and session progression (p<0.0001). However, the lack of a significant interaction between condition and session progression (p=0.5715) suggests that beta-band ERD does not demonstrate enhanced sensitivity to longitudinal improvements in motor imagery proficiency, unlike the mu-band ERD, which showed progressive changes over time. Post hoc comparisons indicated strong initial task engagement on Day 0 (p<0.0001) but showed diminished differentiation between MI-Beta and Rest-Beta in later sessions (p>0.05). This transient nature of beta-band ERD may reflect its broader role in motor control and inhibition rather than specific engagement during motor imagery.^46^ **Figure SI24** shows topographical maps reveal a less focused spatial pattern for beta-band ERD, further suggesting its weaker task specificity.^47–49^

The topographical maps in **Figures 4i and 4j** provide spatial insights into cortical activity during motor imagery (MI) and rest, complementing the temporal trends observed in **Figures 4b,4c,4e, and 4f**. For the mu-band (**Figure 4i and SI23**), motor imagery elicited strong contralateral (C3) desynchronization over the sensorimotor cortex. This finding reflects the specific recruitment of motor-related cortical regions during the mental rehearsal of movement. The minimal ipsilateral (C4) desynchronization observed in the mu-band is in agreement with past studies, which suggest that mu-band activity is predominantly driven by contralateral sensorimotor activity and exhibits only weak bilateral involvement^50^. The rest condition displayed negligible desynchronization, further emphasizing the task-specific modulation of the mu-band during motor imagery. In contrast, the beta-band (18–30 Hz) topographical maps in **Figure 4j** exhibited broader, less spatially distinct patterns of desynchronization. While some contralateral desynchronization over the sensorimotor cortex was evident during MI tasks, it was less pronounced than in the mu-band, consistent with the beta-band’s broader functional role in motor planning, control, and inhibition.^47^ During rest, the beta-band displayed diffuse, modest desynchronization across both hemispheres, indicating lower task specificity. These findings align with the temporal results in **Figures 4h and SI24**, where beta-band ERD was significant during early sessions but showed weaker session-to-session improvement compared to the mu-band.^46^ Overall, the ability of the MindStretcH EEG device to capture these nuanced changes across sessions underscores its suitability for extended BCI applications and its potential for broader applications in neural modulation monitoring.

To assess the signal quality of MindStretcH in comparison to a standard EEG cap, we analyzed trial rejection rates at various amplitude thresholds.^51^ The trial rejection rates, as shown in **Figure 4k**, are comparable between the MindStretcH system and the gold-standard EEG cap with wet conductive Ag/AgCl electrodes (control) across a range of amplitude thresholds (50–150 µV). Statistical analysis with two-way repeated measures ANOVA revealed no significant interaction effects between rejection threshold and device type (p = 0.9979). While there was a significant effect of device type (p = 0.0287), Šídák’s multiple comparisons test did not detect significant differences at any individual threshold level (p>0.05). This suggests that the proportion of “clean trials” relative to all recorded trials of MindStretcH is statistically similar to that of the control. **Figure 4l** presents the root mean square (RMS) amplitude of “clean trials”, which reflects the signal quality of the “clean trials”. The RMS amplitudes for MindStretcH were slightly higher on average than those of the standard EEG cap, but this difference was not statistically significant (p = 0.7922). This indicates that MindStretcH reliably captures neural signals without introducing additional noise or signal distortion, confirming its utility in capturing high-fidelity EEG signals across experimental conditions.

A composite metric was proposed to create an overall signal quality evaluation metric. This metric integrates trial rejection rate and overall RMS signal quality into a single measure. **Figure 4m** shows composite metric scores for the one-time control measurement and the measurements across five sessions (Days 0, 4, 8, 15, and 30) with MindStretcH. As the nature of the composite metric, the lower the values the better overall signal quality. For most subjects that performed recordings on both systems, statistical analysis revealed no significant differences between MindStretcH and the one-time control system measurement at most time points (p > 0.05). However, significant differences between control and MindStretcH were observed at independent time points in subject 3. These results demonstrate that while MindStretcH maintains robust signal quality comparable to the standard EEG cap at most time points, there are some session-specific variations in performance. These differences may reflect factors such as variability in electrode-skin contact quality, operator consistency during device application, or even subject-specific conditions at certain time points. Importantly, these variations showed no consistent trend across subjects or sessions, indicating that MindStretcH can deliver reliable signal quality over extended recording periods.

**Figure 4n** depicts the scalp-electrode impedance values for MindStretcH across five recording sessions (0, 4, 8, 15, and 30 days). Each data point represents the impedance of one of the 13 channels, accumulated across all subjects. The results demonstrate a gradual reduction in impedance over time, with lower impedance values observed in the later sessions compared to the initial ones. Although the two-way repeated measures ANOVA indicated no significant interaction effects (day0 vs day4: p= 0.0632, day 4 vs day 8: p=0.9361, day 8 vs day 15: p=0.9821, day 15 vs day 30: p=0.7618), post hoc analysis (Tukey’s test) identified significant differences between session 1 and later sessions (day 8, p=0.0060; day 15, p=0.0008; day 30, p<0.0001). This trend likely reflects a combination of factors. As the experiment progressed, the operator likely improved their technique in applying MindStretcH, achieving more consistent alignment and pressure distribution across all 13 channels. This improved consistency could lead to better electrode-skin contact, reducing impedance values. Additionally, the hydrogel electrodes were rinsed with water between sessions to remove residual dirt and gel. This cleaning routine might have softened the electrode tips, allowing them to better conform to the scalp’s surface during subsequent sessions. Enhanced conformity increases the effective contact area, thereby reducing impedance. Subject-wise impedance is as shown in Figure SI25. Finally, Figure SI26 shows the head circumference and ear-to-ear length of the study participants (n = 6, female: male = 1:1), with mean values of approximately 57.4 cm for circumference and 39.3 cm for ear-to-ear length. The low variance in head size across subjects suggests that both systems were applied under comparable geometric constraints, reducing potential variability in impedance caused by differences in head dimensions. While the overall variance in head dimensions across subjects was relatively low, the gold-standard EEG cap failed to be worn by the subject with the largest head circumference. This limitation highlights a significant challenge with traditional EEG cap designs, which often lack enough adaptability to accommodate diverse anthropometric characteristics. In contrast, MindStretcH successfully fit all participants, demonstrating its adaptability to a wide range of head sizes and hair conditions. This adaptability underscores the advantages of MindStretcH’s kirigami-inspired mesh design and the application of EGaIn, which ensures compliance with the users even under variable geometric constraints, making it particularly well-suited for diverse populations and real-world applications.

### Machine Learning-Driven Multi-Week Closed-Loop Motor Imagery-based BCI Operation

Building upon the successful demonstration of MindstretcH’s long-term stability and effectiveness in offline EEG recordings, we sought to validate its performance in a more challenging real-world scenario: an online closed-loop BCI system. This step is crucial for assessing MindstretcH’s potential in practical BCI applications that require continuous, real-time interaction between the user’s brain activity and the external device. The subject group was identical to the offline MI EEG recordings. All online experiments were conducted following the completion of the one-month longitudinal offline recordings. **Figures 5a-f** illustrate the machine learning-driven closed-loop motor imagery-based BCI framework, including EEG acquisition, preprocessing, covariance matrix computation, Riemannian geometry-based classification, and real-time feedback. Detailed descriptions of these processes are provided in the “Methods” section. The performance metrics for the online BCI experiment are presented in **Figures 5g–i**, with subject-specific results presented in **Figures SI27 and SI28**.

**Figure 5.**
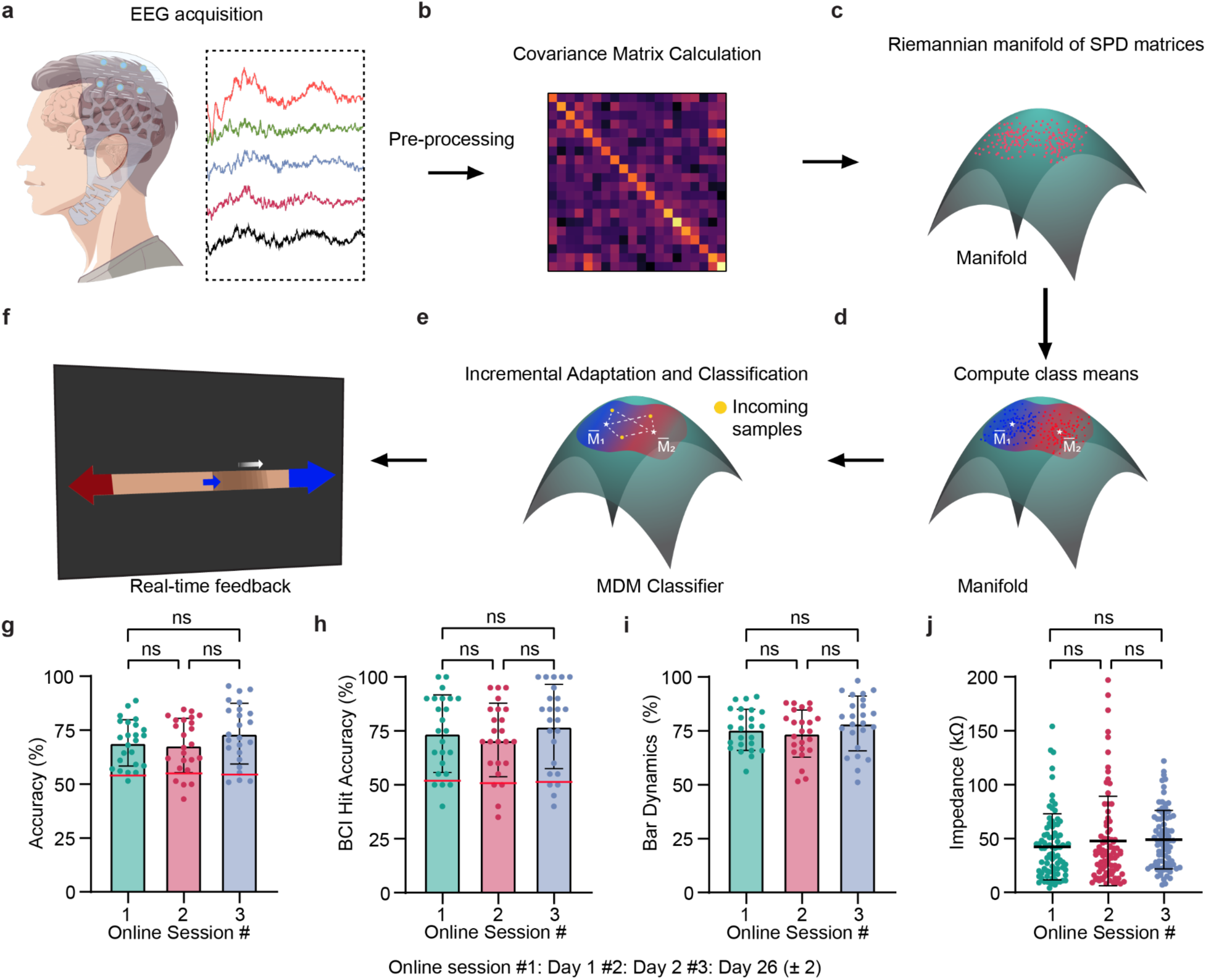
Overview of the online BCI experiment and performance metrics using the MindStretcH device across sessions over time. **(a)** EEG signals were recorded during motor imagery (MI) tasks using a multi-channel setup. (**b**) Preprocessed EEG data were used to compute covariance matrices, which were normalized to reside on the Riemannian manifold of symmetric positive definite (SPD) matrices depicted in (**c). (d)** Class prototypes were computed as averages of the covariance matrices for the MI and Rest classes. (**e**) Incoming samples were incrementally adapted using generic recentering approach and classified based on a minimum-distance-to-mean decoder. (**f**) Participants received real-time feedback via a bar that moved dynamically based on the accumulated evidence from the BCI decoder for the two classes (rest vs. dominant hand MI). (**g–i**) Performance metrics for online BCI sessions 1, 2, and 3 over a month period: (**g**) Sample-wise classification accuracy, **(h**) BCI hit accuracy, and (**i**) bar dynamics, showing group-level averages, individual subject data, and chance level (horizontal red line) (**j**) Impedance levels recorded during sessions 1, 2 and 3, showing consistent interface quality over time.

To evaluate the performance of the online brain-computer interface (BCI), three distinct metrics were employed: sample-wise classification accuracy, bar dynamics, and BCI hits accuracy. Sample-wise classification accuracy measures the percentage of individual EEG samples correctly classified into their respective classes, providing a direct assessment of the classifier’s effectiveness independent of temporal thresholds. In contrast, bar dynamics reflects the user’s control over the BCI system during motor imagery tasks by calculating the proportion of time within a trial that the accumulated evidence supports the correct class, emphasizing sustained user control over time. Meanwhile, BCI hits accuracy evaluates command delivery performance at the trial level by determining the ratio of correct threshold hits to total threshold crossings, normalized to exclude timeout trials. While sample-wise accuracy focuses on granular classification performance, bar dynamics captures temporal control dynamics during task execution, and BCI hits accuracy assesses system-level command reliability across sessions. Together, these metrics offer a comprehensive evaluation framework for understanding classifier precision, user interaction quality, and system-level command delivery in online BCI applications.

**Figure 5g** shows sample-wise classification accuracy across three online sessions. The repeated measures ANOVA revealed no significant differences between sessions (p = 0.1226), indicating stable classification performance over time. The group-level averages were approximately 69% for session 1 (day 1), 68% for session 2 (day 2), and 73% for session 3 (day 26 ± 2). This indicates that the classifier reliably distinguished between rest and dominant hand MI classes. Subject-specific results (**Figure SI27**) highlight variability among participants, with certain participants demonstrating improved classification accuracy along the experiment’s progress. For instance, Subject 2 exhibited significant improvements in classification accuracy from session 1 to session 3 (p < 0.001), while Subject 4 showed significant increases between session 2 and session 3 (p < 0.01). These findings aligned with the results reported in prior research that individual learning effects or increased familiarity with the system may contribute to enhanced performance for certain users.^13^ The BCI hit accuracy results are shown in **Figure 5h**, with individual subject data presented in **Figure SI28**. **Figure 5h** reflects the accuracy of the commands delivered during online runs. Group-level averages were approximately 74% for session 1, 71% for session 2, and 77% for session 3. The repeated measures ANOVA indicated no significant differences between sessions (p = 0.2755) over a month. This metric highlights the robustness of the BCI system in translating neural activity into actionable outputs. While individual variability was observed, most participants maintained high levels of performance across all sessions. **Figure 5i** presents the bar dynamics metric results, which show that group-level averages were approximately 76% for session 1, 74% for session 2, and 78% for session 3. The repeated measures ANOVA again showed no significant differences between sessions (p = 0.1957) over a month. This result indicates stable user control over the bar feedback system. Individual trends suggest that some participants improved their control over time, potentially benefiting from increased familiarity with the task and feedback mechanisms. **Figure 5j and Figure SI30** illustrate impedance measurements across three online sessions as a measure of electrode-skin contact quality and overall signal acquisition stability for the recording sessions. **Figure 5j** presents group-level averages consistent across sessions: 46.3 kΩ for all three sessions (session 1: 42.2 ± 30.6 kΩ, session 2: 47.7 ± 41.5 kΩ, session 3: 48.9 ± 27.1 kΩ). The repeated measures ANOVA revealed no significant differences between sessions over a month (p =0.3938).

These results collectively demonstrate that our MindStrecH system, together with the applied Riemannian geometry-based Minimum Distance to Mean (MDM) classifier^13,40,52^ provides robust and consistent performance over a period of a month at the group level. The stable classification accuracy, BCI hit accuracy, and electrode-skin impedance highlight the reliability of the system. Similarly, the bar dynamics metric further underscores the effectiveness of user control in distinguishing between rest and dominant hand MI classes under real-time conditions. Individual variability observed in all three metrics suggests that user-specific factors play a role in performance outcomes. For some participants, improvements between sessions may reflect learning effects or increased familiarity with the BCI system. These findings emphasize the importance of designing adaptive systems to accommodate individual differences to optimize performance. The consistency of group-level results across sessions also suggests that the system is resilient to temporal non-stationarities in EEG signals. This robustness is likely supported not only by the stability of MindStrecH to reduce the hardware non-stationarity but also by the Generic Recentering (GR) mechanism, which updates class prototypes after each run to account for variability in neural activity patterns.

## Discussions

In this study, we developed the MindStretcH system, which, through its kirigami-inspired mesh structure, EGaIn interconnects, and innovative hydrogel conical electrode platform, achieves efficient adaptation to diverse head sizes and complex scalp geometries while ensuring high-fidelity signal transmission under dynamic conditions. Compared to other research such as that by Tian et al.,^23^ our system demonstrates substantial advantages in hair compatibility, localized pressure distribution, and long-term EEG monitoring stability—particularly excelling in handling dense hair environments and varied anthropometric parameters. Nonetheless, we acknowledge some minor limitations in our work. First, although finite element analysis and extensive mechanical tests have validated the system’s stability under various dynamic deformation conditions, the device may still experience slight performance fluctuations due to external factors such as ambient temperature and humidity during long-term use. This aspect requires further validation and optimization in continuous wear and large-scale applications. Secondly, although we have attained commendable results in longitudinal EEG recording studies, MindStretcH has not yet been implemented in clinical settings for evaluation under more demanding conditions scenarios. Moreover, our detailed exploration of the hydrogel conical electrode platform has shown significant benefits in mitigating hair interference and reducing localized pressure. However, slight variations in electrode-to-skin contact caused by different operators could introduce variability. We posit that future implementation of automated calibration and intelligent feedback mechanisms will further reduce human-induced variability in signal acquisition. Overall, although certain aspects of our study require further refinement, these minor limitations do not detract from the substantial advantages of our system in terms of adaptability, signal stability, and user comfort. Instead, they provide clear directions for subsequent optimization.

In conclusion, our MindStretcH system represents a transformative advancement in non-invasive EEG technology. By integrating a kirigami-inspired, stretchable design with robust EGaIn interconnects and an innovative, hair-compatible hydrogel conical electrode platform, we have successfully addressed key limitations of conventional EEG systems—namely, adaptability, user comfort, and signal fidelity under dynamic conditions. Looking ahead, we plan to expand the system’s multichannel capabilities and incorporate advanced signal processing algorithms to further enhance its performance and usability. We are excited by the prospect of translating these advances into clinical and home use applications, where next-generation EEG systems can drive breakthroughs in neurorehabilitation, intelligent prosthetic control, and personalized brain–machine interfacing.

## Methods

### Materials

2-acrylamido-2-methyl-1-propanesulfonic acid sodium salt solution (≥98%), N,N -methylenebis(acrylamide) (99%), Potassium chloride (KCl), Ammonium persulfate (APS), and Gallium-Indium eutectic were purchased from Sigma-Aldrich. Glycerol (>99%) was purchased from Thermo Fisher Scientific. All chemicals were used as received without further purification.

### Fabrication of MindStretcH

The fabrication of the MindStretcH device begins with the preparation of a polyester film (McMaster-Carr, USA) as a backing layer (**Figures SI1a**). A 3D-printed MindStretcH substrate mold, designed with a thickness of 4 mm to match the dimensions of female snap connectors, is placed on this backing layer, and Ecoflex 00-30 (Smooth-On, USA) is poured into the mold. A 3D-printed Kirigami mesh template is then inserted into the substrate mold to define the structural layout. Female metal snap connectors (Goochain Technology, China) are positioned within the openings of the Kirigami template, corresponding to the targeted channel locations in the 10-10 international EEG montage (**Figures SI1b)**. Once the Ecoflex cures, the Kirigami template is removed, and the device is flipped to expose its back (**Figures SI1c)**. A polyimide-based laser-cut copper interface connector is attached to the substrate to enable electrical interfacing.

The EGaIn interconnects are then fabricated by stirring EGaIn liquid metal (Sigma-Aldrich, USA) with a magnetic stir bar for 30 minutes to create an EGaIn paste. Using a transfer tape (USCutter, USA) and a customized patterned metal stencil (JLCPCB, China), the EGaIn interconnect layout is printed onto the transfer tape (**Figures SI1d)** and subsequently transferred to the MindStretcH substrate (**Figures SI1e)**. Once transferred, the EGaIn interconnect should be aligned with its corresponding copper connector (**Figures SI1f)**. The copper connectors serve as interfaces for standard pin connectors and other extension components. To encapsulate the device, a 1-mm-thick 3D-printed peripheral mold is vertically aligned onto the substrate mold along with the Kirigami template. An additional 1-mm-thick layer of Ecoflex 00-30 is poured into this assembly and cured (**Figures SI1g)**. Finally, after releasing the device from the molds, GKAmp hydrogel-integrated conical porous electrodes are snapped onto the MindStretcH substrate to complete its assembly (**Figures SI1h and SI2**).

### Kirigami-inspired mesh design

The kirigami-inspired mesh design of the MindStretcH wearable EEG system leverages the effort of a prior study.^29^ This approach allows for precise tuning of mechanical properties through adjustments in unit cell parameters, such as horizontal spacing (L_HS_), vertical spacing (L_VS_), and cut length (L_CL_). In our design, we strategically differentiated the kirigami patterns for the temporal and scalp regions to address their distinct functional requirements. For the temporal region, the pattern features L_CL_ = 20 mm, L_HS_ = 5 mm, and L_VS_ = 2.5 mm, yielding an aspect ratio (L_CL_/L_HS_) of 4.0. For the region on the scalp, the pattern uses L_CL_ = 15 mm, L_HS_ = 10 mm, and L_VS_ = 5 mm, resulting in an aspect ratio of 1.5.

### Finite element analysis (FEA)

FEA simulations of MindStretcH were conducted using Autodesk Fusion 360’s Simulation mode to evaluate the mechanical performance of MindStretcH under tensile loading conditions. Material properties for MindStretcH were set as follows: Young’s modulus = 125 kPa, Poisson’s ratio = 0.49, and ultimate tensile strength = 1.38 MPa. The kirigami-inspired structure was modeled with parabolic solid elements, generating 150,771 nodes and 93,153 elements to ensure computational accuracy. Boundary conditions included fixing both the left and right ends of the device (Uy = Fixed, Ux= Uz = Free) and the top and the bottom faces of the device (Uy = Uz = Fixed, Ux = Free) while allowing controlled displacement to simulate an ear-to-ear distance of 40 cm. The force is applied across multiple regions to match the experimental scenario, with a maximum magnitude of -1.2 N along the x-axis. Outputs analyzed included displacement distribution and stress concentration.

Conical electrodes’ FEA simulations were conducted using COMSOL Multiphysics to evaluate the mechanical performance of porous and non-porous elastic 3D-printed electrode molds under compression. The material properties of the electrodes were set to approximately align with the 3D-printed material, with a Young’s modulus of 2 MPa, a Poisson’s ratio of 0.49, and a density of 1010 kg/m³. The scalp was modeled as a soft tissue with a Young’s modulus of 300 kPa, a Poisson’s ratio of 0.49, and a density of 1100 kg/m³. A linear elastic material model was applied for both materials.

The electrode geometries were meshed using free tetrahedral elements, generating approximately 594,726 degrees of freedom to ensure computational accuracy. Boundary conditions included fixing the base of the electrode molds while allowing the tip to deform freely under compression. A uniform downward displacement was applied to simulate interaction with the scalp under realistic EEG system conditions. Outputs analyzed included von Mises stress distribution and displacement fields on the scalp surface.

### Experimental configurations

#### Electromechanical testing

Electrical response under strain was recorded using the chronoamperometry method via a potentiostat (SP-300, BioLogic, France). All mechanical tests were performed using a custom tensile testing setup adapted from a reported protocol by Hsieh et al.^27,53^ A modified 3D-printed holder secured the base of the interconnect samples, while the opposite end was attached to the Arduino-controlled (UNO REV3, Arduino) motor-driven extension mechanism (ramp rate of 68 mm/min, **Figure SI4**). The potentiostat was connected to the interconnect via the two-electrode method. The working electrode (WE) is connected to one end of the interconnect. A 10 Ω resistor was placed in series with the interconnect to regulate current during testing (**Figure SI5**) and connected to the counter/reference (CE/RE) of the potentiostat. The potentiostat maintained an initial applied voltage of 1 V, and measurements were taken at a sampling frequency of 10 Hz. Strain values were recorded in real-time along with current and electrode voltage.

#### Signal fidelity under deformation experiments

To achieve EEG-mimicking signal levels, the EGaIn interconnect was incorporated into a grounded and shielded voltage divider circuit designed to attenuate the input signal to biologically relevant ranges. (**Figure SI6**) The circuit consisted of 5 Ω and 1 MΩ resistors, forming a voltage divider that reduced a 4V sinusoidal wave input signal from a function generator to a microvolt range (∼20 µV). The interconnect was mounted in a 3D-printed holder, ensuring a stable connection for integration with an EEG amplifier/ recorder (ANT Neuro, Netherlands). The circuit configuration incorporated grounding and shielding techniques to minimize electromagnetic and power-line interference, ensuring fair comparison across temporal conditions across experimental groups.

Signal measurements were recorded across three EEG-representative frequencies: 5 Hz, 15 Hz, and 30 Hz. The interconnects were subjected to controlled mechanical deformations, simulating real-world conditions in wearable EEG applications. The following deformation scenarios were applied (**Figure SI8**): 1) Stretching: The interconnect was incrementally elongated to strain levels of 0%, 25%, 50%, 75%, and 100% with 3D printed modules with designated lengths to add to the 3D printed holder, 2) Bending: The interconnect was bent to angles of 0°, 45°, 90°, 120°, and 160° with 3D printed modules with a designated angle to create the desired angle, and 3) Twisting: The interconnect was twisted to half, full, double, and five full rotations. Each deformation condition was recorded for a 5-second duration at each frequency.

### Signal analysis for electromechanical and signal fidelity tests

#### Electromechanical testing

Hysteresis (H) was evaluated for electromechanical coupling and elastic recovery behavior of EGaIn interconnects. To acquire H, a complete strain cycle (100%) was performed and recorded. The calculation was done using:

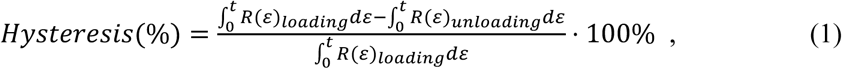

Where ε is strain (%), R_loading_, and R_unloading_ correspond to resistance values during the loading and unloading phases of the strain cycle, respectively.

#### Signal fidelity under deformation

All the signals were collected using LabStreamingLayer.^54^ The collected data of each deformation condition (format and level) and frequency was segmented, post-processed, and analyzed. Specific process and analysis steps include notch filtering (60 Hz), normalization, and segmentation for comparison across conditions (within deformation format).

Attenuation loss calculation: Attenuation loss was calculated for each deformation level as the reduction in signal power from baseline (original signal). This was calculated as follows:

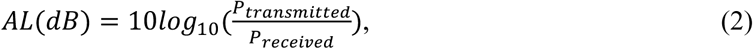

Where *P_transmitted_* represents the baseline input power, and *P_received_* represents signal power received by the amplifier under each deformation. This calculation was repeated for all deformations and frequencies to quantify power attenuation.

Signal-to-Noise Ratio (SNR): SNR was calculated by comparing signals collected by deformed EGaIn interconnects to an ideal baseline, quantifying the impact of deformation on signal integrity. The ideal baseline signal was established by recording the signal generated directly from the signal generator through a medical-grade EEG lead (Bio-Medical, USA) into the amplifier, ensuring an accurate reference for subsequent measurements. SNR is calculated as:

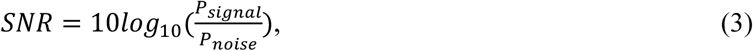

where *P_signal_* represents the recorded signal from the EGaIn interconnect and *P_noise_* represents noise power, which was determined by subtracting the baseline reference signal from the corresponding experimental signals.

#### Synthesis and preparation of GKAmp hydrogel

The hydrogel was synthesized via free radical polymerization. First, sodium 2-acrylamido-2-methylpropane sulfonate (AMPSNa, 4.0 g) and potassium chloride (KCl, 0.1 g) were dissolved in deionized water under continuous stirring at 500 rpm for 1 min. Ammonium persulfate (APS, 0.06 g) was then introduced as the initiator, followed by the addition of N,N′-methylenebisacrylamide (MBAA) at varying concentrations (50, 100, and 150 μL of 70 mg/mL) as the crosslinker. The solution was further stirred at 500 rpm for an additional minute to ensure uniform dispersion. Subsequently, 3.2 g of glycerol was incorporated, and the mixture was continuously stirred under identical conditions. To enhance homogeneity, the solution was subjected to ultrasonic homogenization for 10 s using an ultrasonic processor. Prior to hydrogel film preparation, the pre-polymerized solution was placed in a vacuum desiccator to remove air bubbles and left undisturbed for 5 minutes. The degassed solution was then transferred into designated molds and polymerized in an oven at 60°C for 4 hours. After polymerization, the hydrogel samples were demolded and allowed to cool to room temperature before further characterization. To investigate the effect of potassium chloride concentration on hydrogel properties, a series of GKAmp hydrogels was synthesized by varying the KCl content (0.29, 0.58, and 0.88wt%). The resulting hydrogels were labeled as GKAmpx, where x represents the KCl content relative to the total mass of the hydrogel. For comparison, GKAmp hydrogels without glycerol were also prepared and characterized.

### Mechanical characterization of GKAmp hydrogel

To evaluate the mechanical properties of GKAmp hydrogels, tensile and compression tests were conducted using a custom-built motorized testing platform fabricated via 3D printing (i3 MK3S+, Prusa Research) and integrated with a force gauge (FB200, Torbal). The force gauge has a maximum load capacity of 200 N, and uniaxial strain was applied at a controlled displacement rate of 68 mm/min. Prior to testing, the load cell was calibrated to ensure accurate force measurements. The entire system was regulated by an Arduino-based control unit (UNO REV3, Arduino). For tensile testing, hydrogel samples were prepared with dimensions of 20 mm (width) × 35 mm (length) × 2 mm (thickness), ensuring consistency across experimental evaluations. The effective gauge length of each sample was maintained at 15 mm, with approximately 10 mm of both longitudinal ends securely attached to a non-stretchable adhesive tape to facilitate clamping onto the force gauge and platform. Nominal strain (ε) was defined as the ratio of length change to the original length of the sample. The elastic modulus was determined from the initial linear region (10–15%) of the stress-strain curve by calculating the average slope. For compression testing, cylindrical GKAmp hydrogel specimens with a diameter of 8 mm and a height of 8 mm were utilized. The test was conducted until a terminal strain of 30% was reached. The nominal stress (σ) was computed as the applied force divided by the original cross-sectional area of the sample. To assess the adhesion strength of GKAmp on human skin, a 90° peel test was performed using the previously described motorized testing platform. Each GKAmp sample was fabricated with dimensions of 20 mm (width) × 35 mm (length) × 2 mm (thickness). Before testing, the hydrogel was mounted onto a Kapton film backing layer to provide mechanical support. The sample was then gently pressed onto the forearm of a human volunteer using a 500 g weight for 10 seconds to ensure uniform contact. Each sample underwent a total of 8 attachment-detachment cycles. The adhesion strength was quantified by dividing the maximum shear force by the corresponding sample width. To evaluate the electrical impedance characteristics of GKAmp hydrogels, two complementary measurement techniques were employed. To assess the impedance characteristics of GKAmp hydrogels under varying conditions, an electrochemical workstation (SP-300, BioLogic) was employed. Impedance spectra were recorded over a frequency range of 1–100 kHz. Hydrogel samples were prepared by casting the prepolymer solution into a 3D-printed rectangular mold (25 mm × 25 mm × 3 mm) to ensure uniform sample geometry and volume. To establish electrical connections, two gold bars (10 mm × 20 mm) were embedded at opposite ends of the hydrogel sample, serving as extended probe terminals. These terminals were secured using alligator clips for direct connection to the electrochemical workstation. Ten impedance data sets were recorded per frequency decade, and the corresponding phase angle curves were collected. To investigate the long-term stability of the hydrogel, impedance measurements were repeated weekly over a 4-week period using the same methodology.

### Impedance measurements

First, to further evaluate the skin-electrode impedance performance of GKAmp hydrogels with varying KCl concentrations and benchmark, their impedance was compared against commercial dry and wet electrodes, and an electrochemical workstation (SP-300, BioLogic) was utilized. Impedance spectra were recorded over a frequency range of 1–100 kHz. Hydrogel samples were prepared by casting the prepolymer solution into a 3D-printed rectangular mold (25 mm × 25 mm × 3 mm) to ensure uniform sample geometry and volume. For electrical connection, two gold bars (10 mm × 20 mm) were embedded at opposite ends of the hydrogel sample, serving as extended probe terminals (**Figure SI15**). These terminals were secured using alligator clips for direct connection to the electrochemical workstation. Ten sets of impedance data were recorded per frequency decade, and the corresponding phase angle curves were also collected when applied to human skin. In this setup, GKAmp hydrogels, dry electrodes (OPENBCI Shop), Kendall 200-series foam electrodes (Semi-dry 1), and Kendall disposable surface electrodes H124SG (Semi-dry 2) were affixed to the forearm of a single volunteer as the working electrode. A reference electrode and a counter electrode were placed at distances of 10 cm and 20 cm from the working electrode, respectively. All measurements were conducted on the same subject to ensure consistency. The impedance values obtained were normalized by the electrode-skin contact area to determine the specific skin impedance per unit area. To investigate the long-term stability of the hydrogel, impedance measurements were repeated weekly over a 4-week period using the same methodology.

### Environmental stability assessment

To examine the effect of relative humidity (RH) on hydrogel stability, mass variation of GKAmp hydrogels with and without glycerol was monitored under ambient conditions for 600 hours. The initial mass of each hydrogel sample was recorded immediately after preparation, and subsequent mass measurements were taken at defined time intervals. Mass values were normalized to the initial mass to calculate the relative mass change over time. Concurrently, ambient humidity levels were monitored throughout the duration of the experiment to assess correlations between environmental humidity and hydrogel weight fluctuation.

### Charge injection capacity (CIC) measurements

The electrochemical performance of GKAmp hydrogels was characterized using a three-electrode system to measure charge injection capacity (CIC). The working electrode consisted of a GKAmp hydrogel interface deposited on a platinum (Pt) sheet with dimensions of 1.5 cm × 1.5 cm. A Pt sheet of the same size served as the counter electrode, while an Ag/AgCl electrode immersed in a saturated KCl solution was used as the reference electrode. All electrodes were submerged in a 10 mM phosphate-buffered saline (PBS) electrolyte solution to maintain a consistent electrochemical environment. For CIC measurement, biphasic pulses of ± 0.5 V were applied at an interval of 0.01 s for 1500 consecutive cycles. The total charge injection density (*Q_inj_*) was determined using the following equation:

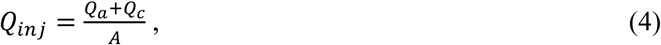

where *Q_a_ and Q_c_* represent the cathodic and anodic charge quantities, respectively, and denote the effective hydrogel-electrode interface area. The CIC values recorded on the first day were normalized, and the long-term electrochemical stability was assessed by repeating the test once per week over a period of four weeks. A total of four independent samples were evaluated in each test to ensure reproducibility.

### Cell viability test

The biocompatibility of the GKAmp was assessed using a cytotoxicity assay based on the extract method. The GKAmp prepared with 3D-printed soft molds was immersed in Dulbecco’s Modified Eagle Medium (DMEM) for 2 hours to obtain the hydrogel extract. A total of 1×10⁵ mouse NIH/3T3 fibroblasts were seeded into a 96-well plate and incubated for 4 hours. Once the cells were fully adhered and proliferating, the culture medium was replaced with the hydrogel extract solution. Cells cultured in standard DMEM medium without hydrogel extract served as the control group. After incubation periods of 24 and 48 hours, cell viability was assessed using the CCK-8 cytotoxicity assay kit following the manufacturer’s protocol. Absorbance measurements were taken at 450 nm using a microplate reader. The relative cell viability was calculated using the following formula:

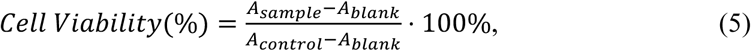

where *A_sample_* represents the absorbance of cells treated with the hydrogel extract, *A_control_* is the absorbance of cells cultured in normal DMEM medium, and *A*_=6-.>_ corresponds to the background absorbance. Experiments were conducted in quadruplicate to ensure statistical significance.

### Human participants and EEG data acquisition

Six able-bodied, healthy participants with normal or corrected-to-normal vision participated in the study (3 females, 3 males, age: 24.3 ± 5.8 y). The experimental protocol was approved by the local ethics commission (STUDY00000516, The University of Texas at Austin, TX, USA). Electrodes were arranged according to the international 10-10 system, with the following channels designated as recording electrodes: FC3, FC4, FCz, C1, C2, C3, C4, C5, Cz, CP3, and CP4. The reference electrode was positioned at CPz. This configuration was chosen to ensure optimal coverage of motor and sensorimotor cortical areas while maintaining high signal quality for motor imagery and rest task analysis. All the signals were collected using LabStreamingLayer (LSL; Swartz Center for Computational Neuroscience, University of California, San Diego, USA) to record in real-time with a 32-channel research-grade EEG amplifier (ANT Neuro, Netherlands). Electrode impedance values were acquired using eego64 software (ANT Neuro, Netherlands).

### One-month longitudinal offline MI EEG recordings

#### Study protocol

Each recording session (days 0, 4, 8, 15, 30) involved participants undergoing four runs of motor imagery (MI) offline recordings using the MindStretcH system. The scalp-electrode impedance measurements were taken at the end of each recording session. Additionally, a single control recording session using a commercial EEG system was conducted for each participant at an arbitrary time point within the one-month experimental period.

Due to the additional preparation time required for the commercial EEG system, each control recording session consisted of 3 runs, whereas MindStretcH sessions consistently included 4 runs per day. This adjustment ensured that the overall duration of each recording session remained comparable across systems, optimizing alignment in experimental conditions and minimizing potential variability in participant fatigue or environmental factors.

#### Interface for offline motor imagery task

The ’bar feedback task’ utilized in the MI experiment is derived from well-established protocols widely adopted in BCI research over the past decade. Variants of this task have been implemented in numerous previous studies.^40,52,55,56^

Each trial begins with a 2-second fixation period, during which participants are presented with a black screen containing white text instructing them to resume focus and prepare for the upcoming task. This is followed by a 2-second cue period, during which the type of task is indicated by the color of the oval icon: a blue oval signals an upcoming rest task, while a red oval signals an upcoming MI task. The cue period transitions into a 6-second task period, during which participants perform either rest or MI tasks as instructed. A running progress bar at the bottom of the screen provides real-time feedback by loading from 0% to 100% over the 6-second duration. Each trial concludes with a 4-second intertrial relaxation period, allowing participants to reset before the subsequent trial.

For rest tasks, a white cross overlays the blue oval during the task period, visually reinforcing that participants should remain relaxed. For MI tasks, an arrow pointing toward the participant’s dominant-hand side overlays the red oval during the task period, prompting participants to mentally rehearse the kinesthetic movements of their dominant hand. The visual feedback provided by this interface ensures that participants remain engaged and accurately follow task instructions throughout each session.

#### EEG signal analysis and visualization

Raw EEG signals were preprocessed using MATLAB. Preprocessing steps included: 1) Spatial Filtering: A Common Average Reference (CAR) spatial filter was applied to reduce noise and enhance signal quality. 2) Notch Filtering: A 60 Hz notch filter was applied to remove powerline noise. 3) Bandpass Filtering: Signals were first filtered with a fourth-order Butterworth filter to 1–50 Hz to get the broadband EEG signal for any early-stage visualization needs. The signals of interest were then later filtered to between 8–13 Hz (mu-band) or 18–30 Hz (beta-band) using a fourth-order Butterworth filter to retain task-relevant frequency bands.

Each “trial” is composed of a “fixation period,” “cue period,” and an assigned “task,” “motor imagery task,” or “rest task.” They were segmented based on event markers. All trials exceeding a maximum allowed amplitude threshold of 50 µV during the examined period (last 2 seconds from fixation to the end of task) were rejected to minimize contamination from artifacts such as muscle activity or eye movements. Valid trials were retained for further analysis. For each valid trial, feature extraction was conducted to isolate and analyze the mu (8–13 Hz) and beta (18–30 Hz) frequency bands. Bandpass filters were applied to the preprocessed EEG signals to extract these frequency components. To quantify event-related desynchronization (ERD), the power values during the task period were normalized against baseline periods corresponding to the fixation period. Next, grand averages of normalized power in the mu and beta bands were computed across trials for MI and rest conditions. Topographical maps of ERD in the mu and beta bands were created using MATLAB’s EEGLAB toolbox. Data from all participants were averaged for each condition (MI vs. rest) and plotted across electrode locations using standardized channel coordinates. Statistical comparisons between MI and rest conditions at different time points (Day 0, Day 4, Day 8, Day 15, Day 30) were performed using statistical methods described in the following statistical method sections.

### Statistical methods used in EEG analysis

A two-way repeated-measures analysis of variance (RM-ANOVA) was employed to evaluate the longitudinal changes in motor imagery (MI) and rest-related EEG activity in the mu (8–13 Hz) and beta (18–30 Hz) frequency bands. The analysis considered two factors: task condition (MI vs. rest) and time (Day 0, Day 4, Day 8, Day 15, Day 30). The assumption of sphericity was tested and corrected using the Geisser-Greenhouse epsilon where necessary. Statistical significance was set at an alpha level of 0.05. The RM-ANOVA results identified significant effects for the column factor (task conditions) and interactions between task condition and time, indicating distinct differences in normalized power between MI and rest conditions across sessions. Post-hoc comparisons were conducted using Šídák’s multiple comparisons test to assess pairwise differences between conditions at each time point and across time within each condition. Adjusted p-values were reported to control for family-wise error rates.

### Signal quality benchmarking

Signal quality metrics were compared between two systems: the experimental system (MindStretcH) and a control system (a commercial stretchable head cap with Ag/AgCl conductive gel electrodes). For both systems, trial rejection rates, RMS amplitudes, and composite metrics were evaluated across all rejection thresholds to assess noise levels and artifact tolerance.

### Composite metric calculation

To evaluate overall signal quality within each session, a composite metric was calculated for each run as:

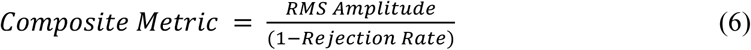

This metric integrates Signal quality (RMS amplitude) with the rate for high signal quality trials (1−rejection rate), where lower values indicate better signal quality. All four runs from each session were analyzed for MindStretcH to calculate composite metrics channel-wise across five rejection thresholds (50 µV to 150 µV). To accommodate the sample size of the control system, the three runs with the lowest average composite metric values were selected for further analysis of direct comparison with the control system.

### Trial rejection rate and RMS amplitude calculations

To evaluate signal quality, a range of rejection thresholds (50 µV, 75 µV, 100 µV, 125 µV, and 150 µV) was applied to identify and exclude trials with excessive amplitude deviations. For each threshold, the maximum absolute value of each trial was calculated and compared with the threshold. RMS amplitudes were calculated for both MI and rest periods across all channels. Rejection rates were computed as the proportion of excluded epochs relative to the total number of epochs.

### Four-week online BCI experiment

After all participants had completed the one-month longitudinal offline MI EEG recordings, this multi-week online BCI experiment commenced. Participants engaged in a motor imagery (MI) task to control a one-dimensional (1D) bar feedback system. In contrast to the one-month offline MI EEG recordings, this experiment involved distinguishing MI of the dominant hand from rest using a subject-specific Riemannian geometry-based Minimum Distance to Mean (MDM) classifier to provide real-time feedback and BCI commands. Prior to the first online BCI session, participants completed four runs of offline calibration on the same day. Each run consisted of multiple trials alternating between MI of the dominant hand and rest. The data collected during these runs was used to train the Riemannian MDM decoder.

The online experiment was conducted across three sessions:

- First Online Session: Participants performed four online runs immediately following the offline calibration on the same day.
- Second Online Session: This session was conducted on the second day of the experiment, which had four online runs.
- Third Online Session: The final session used the same procedure as the second one and took place approximately 26 days later (± 2 days), allowing for a four-week separation from the initial sessions.

To optimize user performance and ensure familiarity with the system, participants completed one practice run and one threshold-tuning run prior to each set of four online runs. These preparatory runs provided an opportunity for participants to adapt to the system and allowed for adjustments to thresholds, ensuring a tailored and effective user experience during the experiments.

### Feature extraction

Signals were bandpass filtered with a second-order Butterworth filter in [8, 30] Hz to capture sensorimotor rhythms (SMRs). The filtered EEG data sample matrix, denoted as *X* ∈ *R^Nc^*^×C^*^t^*, represents signals from *N_c_* channels over *N_t_* time points. Normalised sample level covariance matrices, *M_i_*, were computed for each sliding 1-second window using a shrinkage-based estimator (Ledoit and Wolf method^57^) to enhance numerical stability:

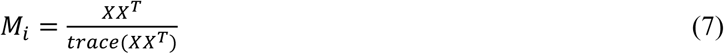

### Riemannian geometry framework

Covariance matrices derived from EEG signals are treated as points on a Riemannian manifold, which is a smooth, differentiable space where each point has an associated tangent space that is Euclidean.^58^ The parametric equation of the shortest path, or geodesic, for curved spaces between two matrices *M*_1_ and *M*_2_ on the Riemannian manifold can be expressed parametrically as:

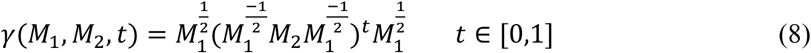

Using the parametric equation (8) the riemannian distance between two covariance matrices *M*_1_ and *M*_2_ can be computed as:

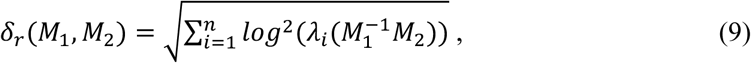

where λ_*i*_ are the eigenvalues of *M*_1_-^1^*M*_2_. The Riemannian distance in equation (9) also possesses the affine invariance property and is commonly referred as Affine-Invariant Riemannian Metric (AIRM).^59^ Using the above mentioned definitions of geodesics and distances, riemannian mean for a set of covariance matrices could be estimated by solving the optimization problem^60^ in equation (10)

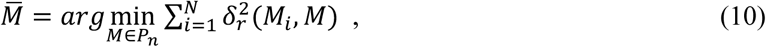

where *P_n_* is the space of SPD matrices, and *N* is the number of covariance matrices and δ*_r_* is the riemannian distance.

The Minimum Distance to Mean (MDM) classifier is one of the most widely used models within the Riemannian geometry framework for classifying covariance matrices, and it has demonstrated robustness in online BCIs^13,40,61,62^. MDM is a distance-based nearest neighbor classifier defined on the manifold of symmetric positive definite (SPD) matrices. Specifically, it utilizes class-specific Riemannian means (also referred to as class prototypes) to construct a nearest neighbor classifier. During the calibration phase, Riemannian means are estimated for *k^th^* class (<u>*M_k_*</u>) through (10) using the set of corresponding class-specific covariance matrices. In the online phase, each incoming covariance matrix (*M_test_*) is classified by computing its Riemannian distance, as defined in equation (9), to all class prototypes estimated during calibration, and assigning it to the nearest one (see equation (11))

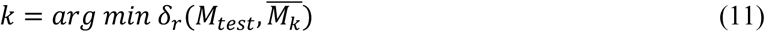

### Online BCI Operation

#### Brain-computer Interface (UI)

In this task, participants received visual feedback via a bar displayed on the screen, which moved horizontally to indicate the classification output. Depending on the participants, the dominant hand side of the bar corresponded to the MI task, while the other side represented the rest task. Each trial began with a 2-second relaxing period (±1 second) to allow participants to settle, followed by a 2-second fixation period. A task cue was then presented for 2 seconds: a blue arrow pointing at the dominant hand side signaled a MI task trial, while a red arrow pointing at the other side indicated a rest task trial.

During offline runs, participants performed either rest or MI as instructed by the task cue. The bar movement during these runs was simulated based on the task cue, providing continuous visual feedback for 5 seconds. In online runs, however, the bar’s movement was controlled in real time by the BCI decoder, which processed EEG data to classify each sample as either rest or dominant hand MI. The bar moved proportionally based on the accumulated evidence for each class. Decision thresholds were displayed as lines on either side of the screen, and if the bar crossed one of these thresholds, discrete feedback in the form of an arrow appeared for 2 seconds to confirm the executed command. If no threshold was reached within 7 seconds, the trial ended with a ‘Timeout.’

Participants were instructed to perform their tasks as follows: “For rest trials, remain still and relaxed without engaging in any physical or mental activity. For dominant hand MI trials, vividly imagine performing a single sustained movement with your dominant hand without physically moving or contracting any muscles. Strive for consistent mental imagery across all sessions.”

#### Real-Time Classification

During online operation, EEG signals were processed in real time to enable dynamic classification. The incoming signals were bandpass filtered using a second-order Butterworth filter in the frequency range of [8, 30] Hz to isolate sensorimotor rhythms (SMRs). Covariance matrices were computed from 1-second sliding windows of the filtered EEG data, updated every 62.5 ms. Class means were computed by averaging the covariance matrices over the Riemannian manifold for the MI and Rest classes, and incoming samples were classified using the Minimum Distance to Mean (MDM) decoder. The classification output was used to provide real-time feedback via a one-dimensional (1D) bar that moved dynamically based on the accumulated evidence for each class.

#### BCI controller

The BCI system estimated the probability distribution of each EEG sample belonging to the respective task classes (e.g., motor imagery vs. rest). To determine a command, the system employed an evidence accumulation approach over time. Specifically:

- A 1-second sliding window of EEG signals was processed every 62.5 ms.
- Probabilities for each class were accumulated until one class’s evidence exceeded a predefined threshold, triggering a command.
- If no threshold was reached within a specified timeout period, the trial was classified as “Timeout” and no command was issued.

To ensure smooth evidence accumulation, an exponential smoothing method was applied:

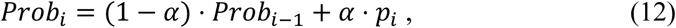

where *Prob_i_* is the accumulated probability for a given class at the *i*-th sample, and *p_i_* is the posterior probability of that class for the *i*-th sample. α is the smoothing factor, which we kept at α = 0.05 to balance responsiveness and stability in real-time feedback.

#### Incremental Adaptation via Generic Recentering (GR)

To account for intra-subject variability and temporal non-stationarities in EEG signals, an unsupervised recentering mechanism was implemented during each online run following the methods reported in reference.^13^ The core principle of this recentering-based domain adaptation approach is to align the mean of the covariance distributions from both the calibration and online sessions to the identity matrix on the Riemannian manifold. Specifically, the mean covariance matrix from the calibration session is computed and used to transform the calibration data via an Affine-Invariant Riemannian Metric (AIRM)-based whitening transformation^63^ prior to training the MDM classifier. During the online session, as a new covariance matrix becomes available, the mean of the set of online covariance matrices is approximately estimated in an incremental fashion (see Kumar et al.^13^ for details). The incoming covariance matrix is then transformed using the same AIRM-based whitening procedure before being classified by the MDM model trained on the transformed calibration data. This incremental recentering procedure is hypothesized to mitigate the distributional shift between calibration and online data and is shown to facilitate robust classification under non-stationary conditions.

### Online BCI performance analysis

To evaluate the performance of the online brain-computer interface (BCI), three key metrics were used to assess classification accuracy, user control levels, and command delivery performance. These metrics are detailed below:

### Sample-Wise Classification Accuracy

This metric measures the percentage of individual samples correctly classified as belonging to the respective class (e.g., MI of the dominant hand vs. rest) during each run. The classification accuracy was calculated as:

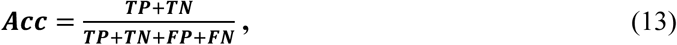

where ***TP, TN, FP*** and ***FN*** correspond respectively to true positives, true negatives, false positives, and false negatives. Unlike trial-level performance, sample-wise accuracy is independent of thresholds used for evidence accumulation, providing a direct measure of classifier effectiveness.

### Bar Dynamics

The bar dynamics metric reflects the user’s level of control over the BCI system during motor imagery (MI) task execution. It represents the proportion of time during a trial that the accumulated evidence from the BCI decoder supported the correct class. Bar dynamics were computed based on the percentage of samples where the accumulated probability for the correct class exceeded 50%, indicating successful control.

### BCI Hits Accuracy

This metric evaluates command delivery performance by calculating the ratio of trials with correct threshold hits to the total number of threshold crossings within a session. Trials resulting in a “Timeout” outcome were excluded from this calculation. The metric was normalized to account for timeout trials using:

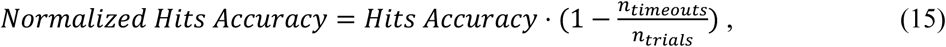

where *n_timeouts_* is number of trials that ended with a timeout and *n_trials_* is total number of trials in a session.

### Chance Levels

To establish baseline performance levels, chance levels for classification accuracy and command delivery metrics were computed by randomly permuting testing labels 10,000 times. The outcomes were averaged over these permutations to determine the expected performance under random conditions.

## Acknowledgements

We would like to thank Nanshu Lu for her guidance and for providing access to the use of the LPKF Protolaser U4 system. H.W. would like to acknowledge support from the UT Austin Startup fund, Alzheimer’s Association New to the Field (AARG-NTF) research grant, Department of Defense (DoD) Defense Advanced Research Projects Agency (DARPA) REM-REST grant, and UT Austin Proof of Concept Award. J-C.H. would like to acknowledge support from the Texas Innovation Center at the University of Texas at Austin. We acknowledge BioRender.com for the figure drawing.

## Author Contribution

Conceptualization, J.H. M.Y., H.W.; Methodology, J.H., M.Y., H.A., S.K.; Software, J.H., H.A., S.K. J.d.R.M.; Validation, J.H., M.Y.; Formal analysis, J.H., M.Y., V.K., H.A. H.D.; Investigation, J.H., M.Y., V.K., Z.A.,; Data curation, J.H., M.Y.; Writing – original draft, J.H., M.Y., H.A., V.K., S.K. J.d.R.M. H.W.; Writing – review & editing, J.H., M.Y., H.A., V.K., S.K., K.W.K.T., W.W., J.J., H.D., T.C., Z.A., D.W., T.E., R.W., A.G., W.H., W.D.M., A.G., J.d.R.M, H.W.; Visualization, J.H., M.Y., H.A., V.K.; Project Administration, J.H., H.W.; Resources, J.d.R.M., H.W.; Funding acquisition, H.W.;

