## Supplementary Information for "Stretchable, hair-compatible, and long-term stable wearable EEG system"

**Figures related to:**
**Design and Performance of MindStretcH**

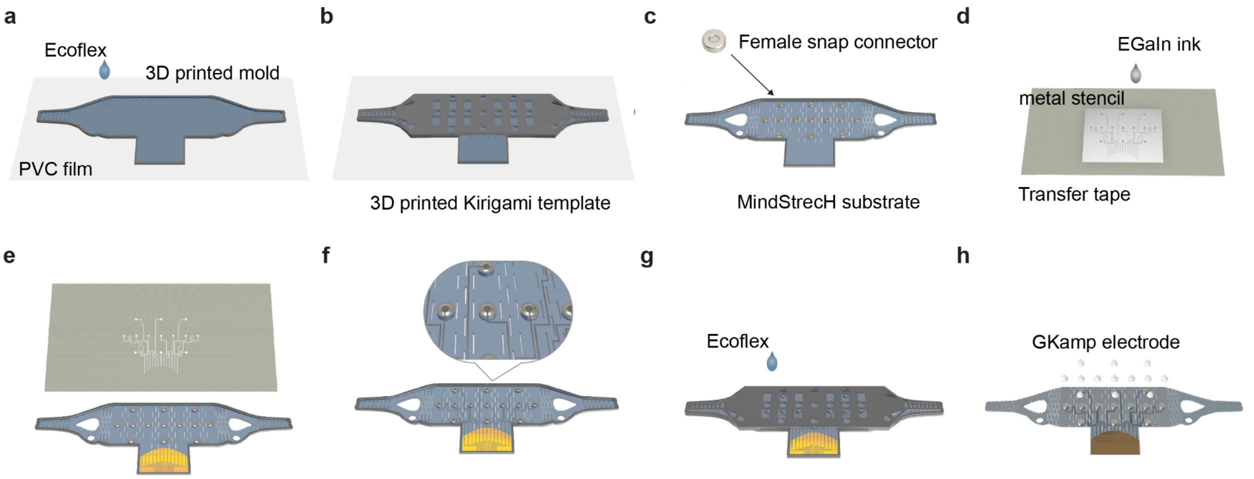


**Figure SI1. Fabrication process of the MindStretcH EEG system.**
(a) Preparation of the substrate using a 3D-printed mold and Ecoflex 00-30 poured onto a polyester backing film. (b) Integration of a 3D-printed Kirigami mesh template and placement of female snap connectors aligned with the 10-10 EEG montage channel locations. (c) Cured MindStretcH substrate after removal of the Kirigami template, flipped to expose the back side for further processing. (d) Application of EGaIn interconnects using a patterned metal stencil and transfer tape to define the conductive pathways. (e) Alignment and attachment of EGaIn interconnect to copper interface connectors on a polyimide film and the substrate. (f) Close-up view showing a precise alignment of EGaIn interconnects with copper connectors for reliable electrical interfacing. (g) Encapsulation of the device with an additional layer of Ecoflex using a peripheral mold for mechanical stability. (h) Final assembly with GKAmp hydrogel-integrated conical porous electrodes snapped onto the substrate, completing the MindStretcH system.

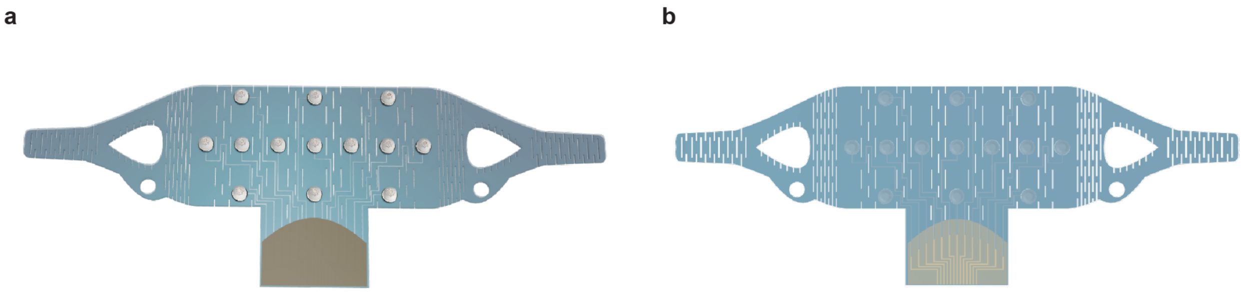


**Figure SI2. Completed MindStretcH EEG device.**

(a) Front view of the fabricated MindStretcH EEG system showcasing its Kirigami-inspired mesh structure and integrated electrodes. (b) The back view highlights the stretchable interconnects and encapsulated components, ensuring durability and adaptability to diverse head morphologies.

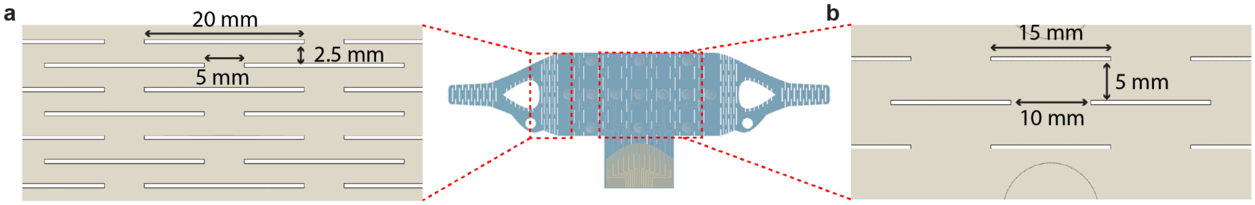


**Figure SI3. Kirigami patterns used in the MindStretcH EEG system.**
(a) Kirigami pattern designed for setting on the temporal region, featuring a horizontal spacing (LHS) of 5 mm, vertical spacing (LVS) of 2.5 mm, and cut length (LCL) of 20 mm, yielding an aspect ratio of 4.0. (b) Kirigami pattern designed for the scalp region, with LHS = 10 mm, LVS = 5 mm, and LCL = 15 mm, resulting in an aspect ratio of 1.5.

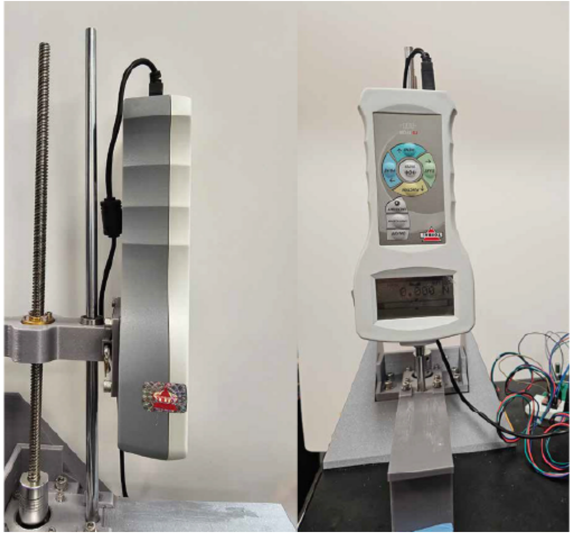

**Figure SI4. Experimental setup for electrical and mechanical characterization of EGaIn interconnects.**
Photographs of the tensile testing system used to evaluate the resistance stability of EGaIn interconnects under cyclic stretching test.

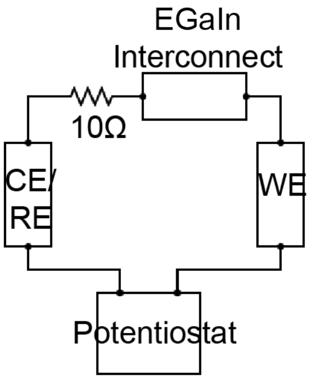


**Figure SI5. Circuit diagram for resistance measurement of EGaIn interconnects.**
Schematic representation of the electrical setup used for resistance characterization, featuring a potentiostat connected to the EGaIn interconnects with a 10 Ω resistor in series. The working electrode (WE), counter electrode (CE), and reference electrode (RE) are designated for accurate resistance monitoring.

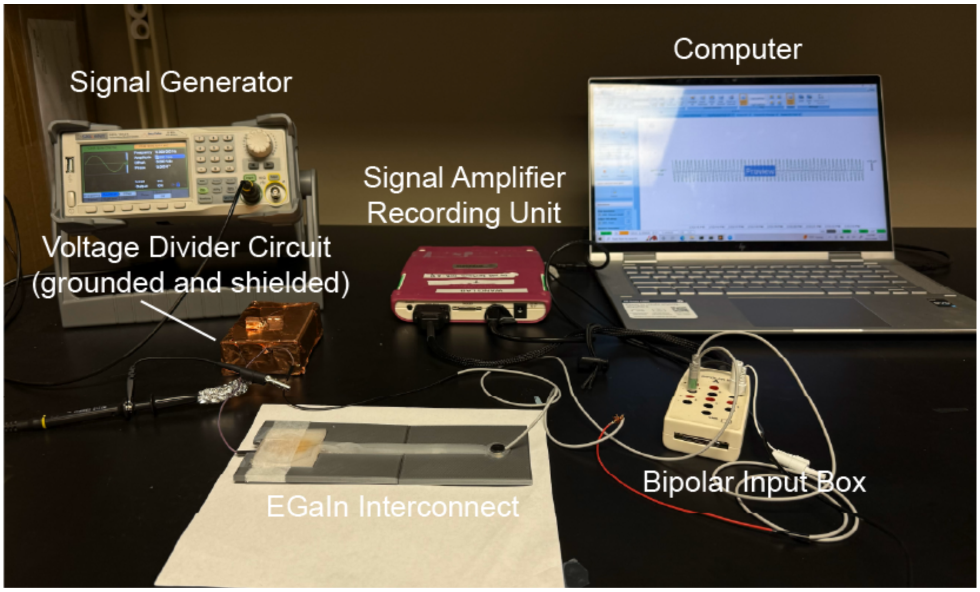


**Figure SI6. Signal fidelity testing setup for EGaIn interconnects.**
Photograph of the experimental arrangement used to assess signal transmission quality through EGaIn interconnects. The setup includes a signal generator, voltage divider circuit (grounded and shielded), signal amplifier/recording unit, and a computer interface for data acquisition and analysis.

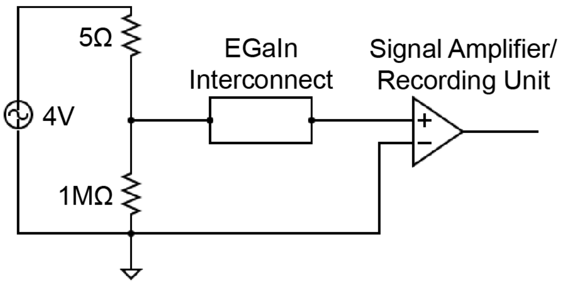


**Figure SI7. Circuit diagram for signal fidelity testing of EGaIn interconnects.**
Schematic of the electrical configuration used to evaluate signal consistency under dynamic strain conditions. The circuit includes a 4 V power source, resistors (5 Ω and 1 MΩ), and a signal amplifier connected to the EGaIn interconnects for real-time monitoring of current flow and signal stability.

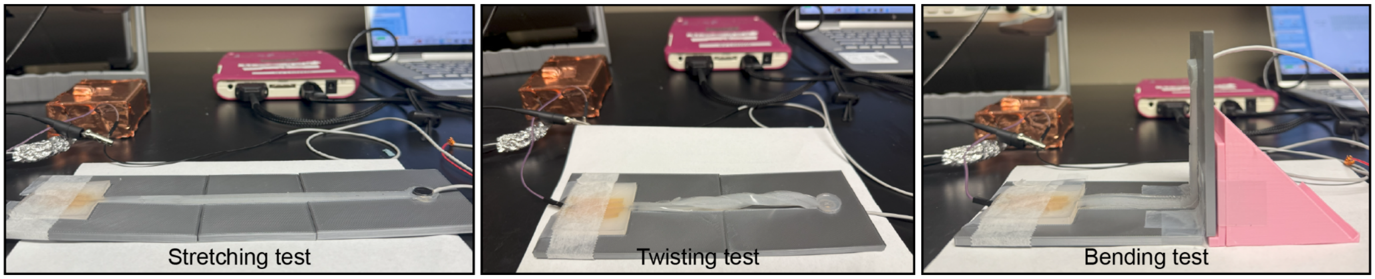


**Figure SI8. Deformation tests for EGaIn interconnect under stretching, twisting, and bending conditions.**

Photographs of the experimental setup used to evaluate the mechanical resilience of EGaIn interconnects under three deformation scenarios: (a) Stretching test with incremental strain levels (0%, 25%, 50%, 75%, and 100%) applied using modular 3D-printed holders. (b) Twisting test with interconnects subjected to varying rotations (half, full, double, and five full rotations). (c) Bending test performed at angles of 0°, 45°, 90°, 120°, and 160° using 3D-printed modules to achieve precise angular deformation.

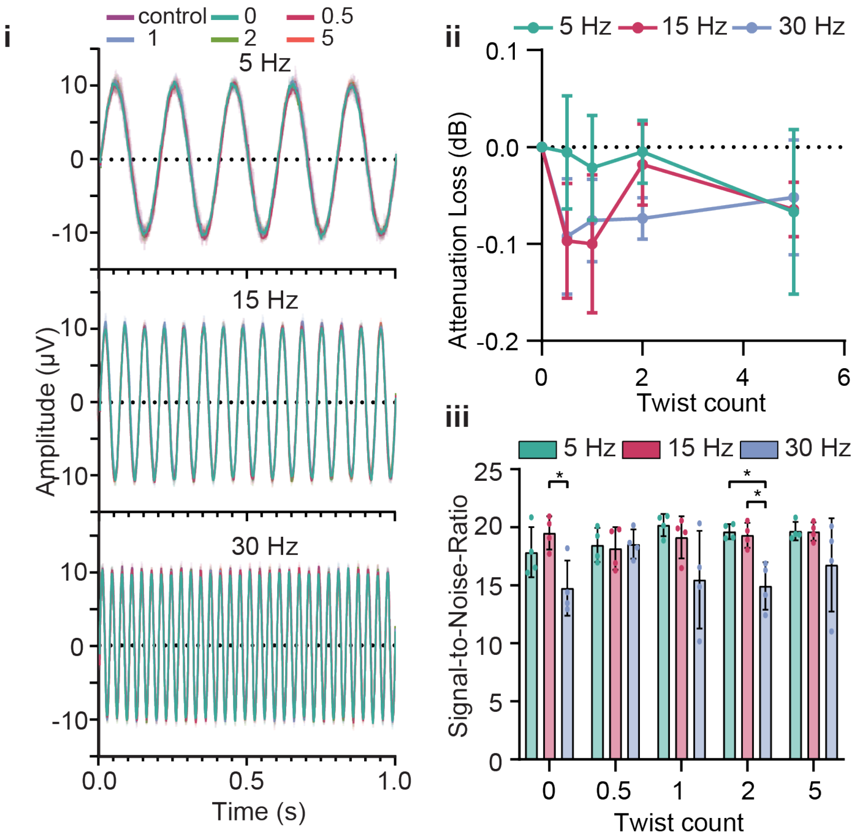


**Figure SI9. Signal fidelity under twisting deformation.**

(a) Overlaid waveforms of EEG-representative frequencies (5 Hz, 15 Hz, and 30 Hz) under different twist counts (control, 0.5, 1, 2, and 5 full twists), demonstrating consistent amplitude and phase alignment. (b) Attenuation loss across twist counts for representative frequencies, showing minimal attenuation (<0.2 dB) across all conditions. (c) Signal-to-noise ratio (SNR) for each frequency under increasing twist counts, consistently exceeding 15.

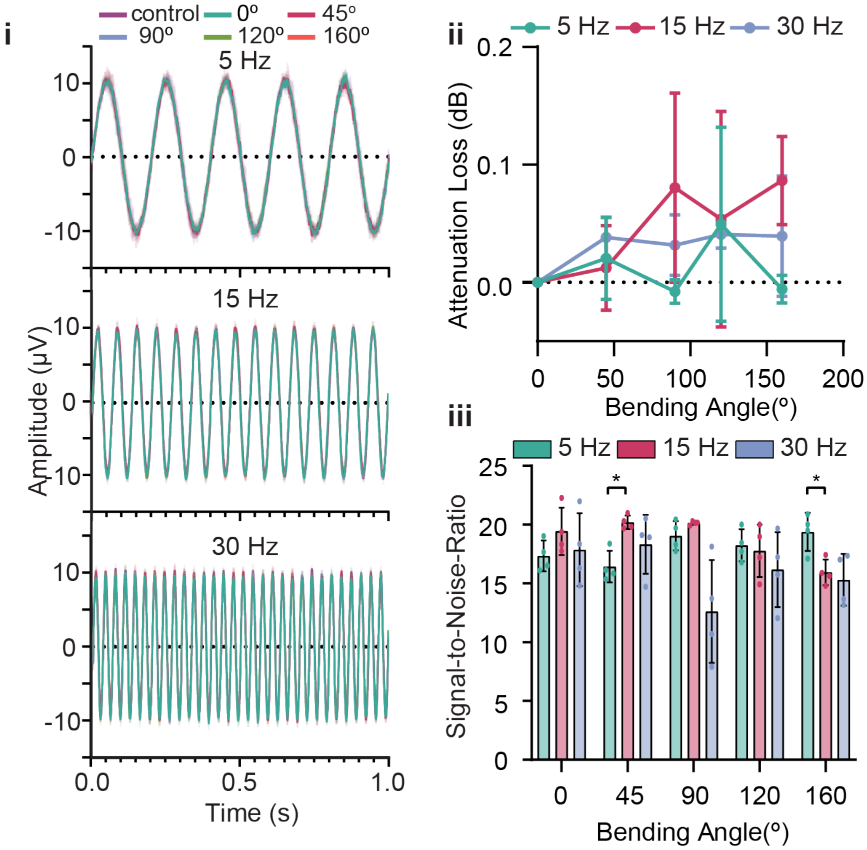


**Figure SI10. Signal fidelity under bending deformation.**

(a) Overlaid waveforms of EEG-representative frequencies (5 Hz, 15 Hz, and 30 Hz) under bending angles (control, 0°, 45°, 90°, 120°, and 160°), showing preserved signal amplitude and phase alignment. (b) Attenuation loss at various bending angles for representative frequencies, remaining below <0.2 dB even at extreme angles. (c) SNR values for each frequency across bending angles are consistently above 15.

**Figures related to: Integration of High-Performance Hydrogel and 3D-Printed Elastic Electrodes for Hair-Compatible EEG Sensing**

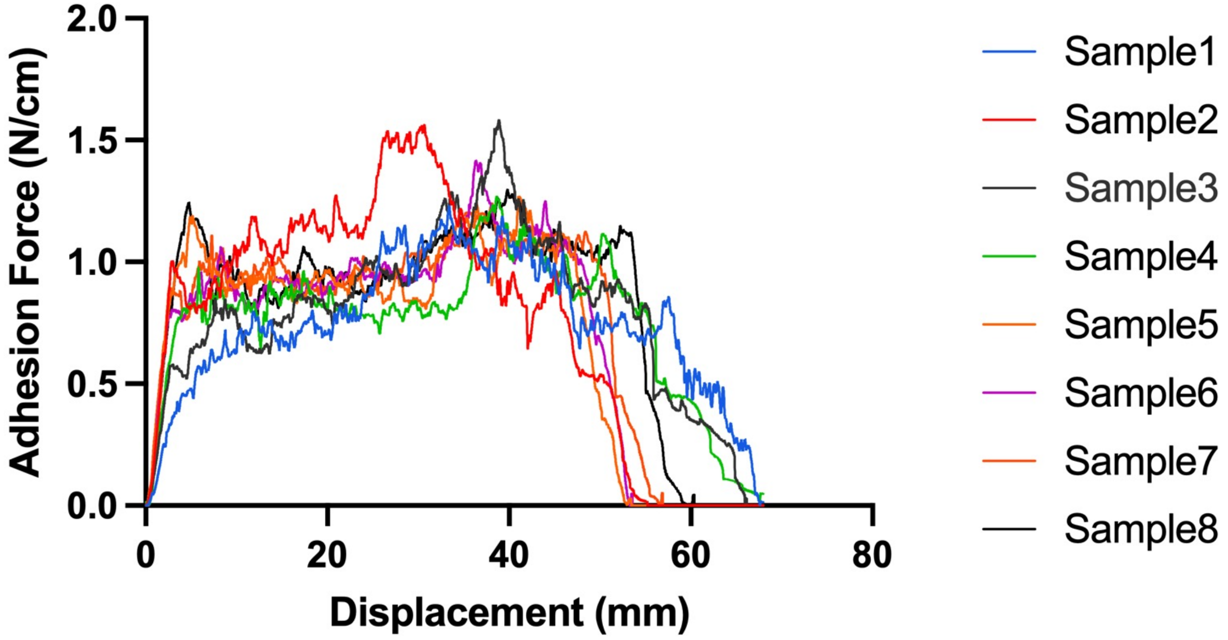


**Figure SI11.** Adhesion force-displacement curves of GKAmp hydrogel after eight repeated cycles of attachment/detachment testing on the skin surface

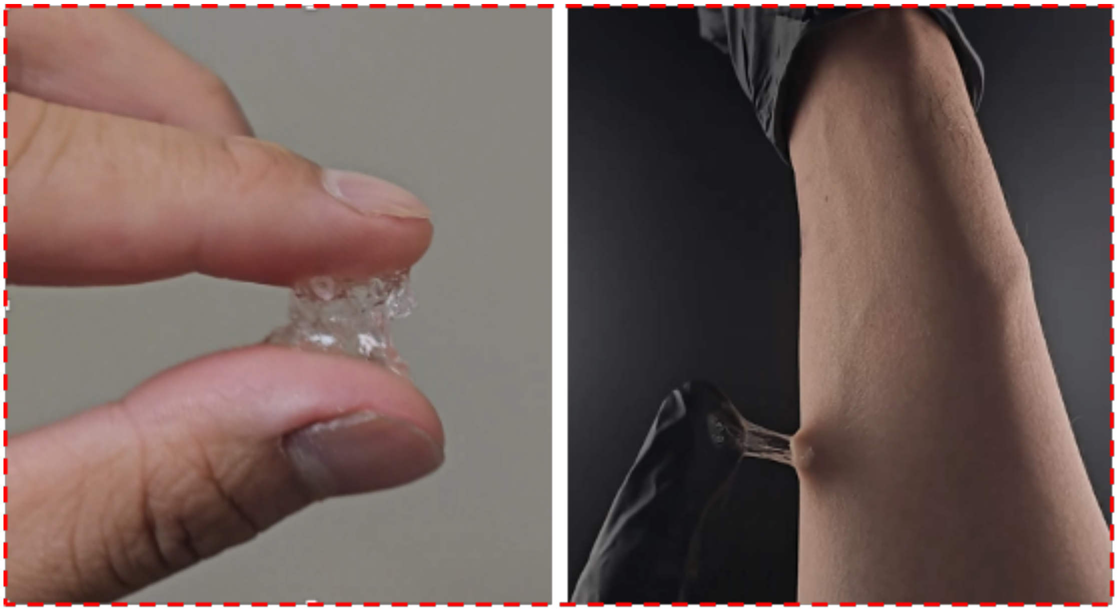

**Figure SI12.** Optical photos of GKAmp hydrogel adhesion on fingers and arms.

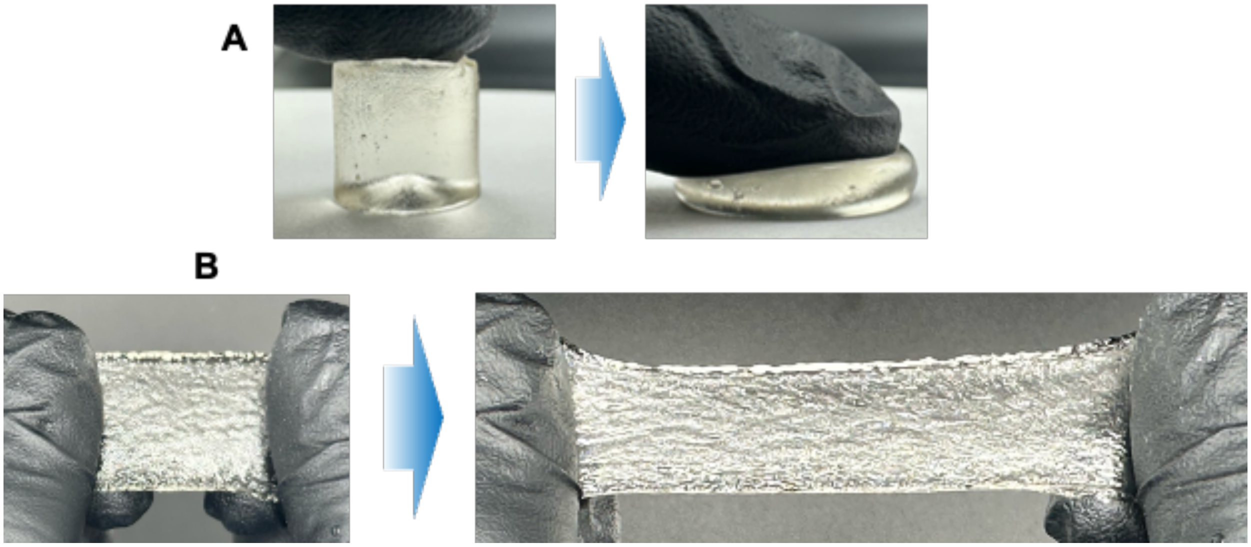


**Figure SI13.** Optical photos of GKAmp hydrogel during compression (top) and stretching (bottom).

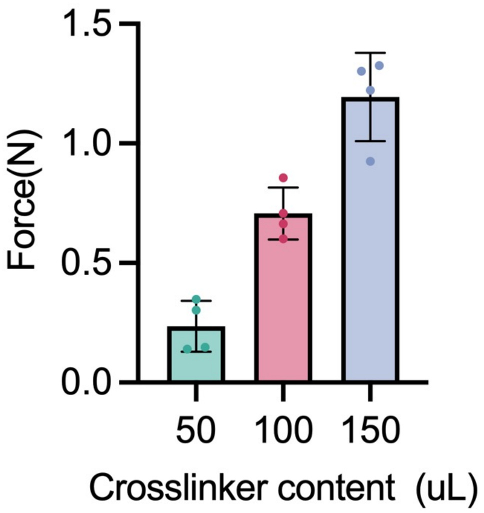


**Figure SI14.** Compressive stress of GKAmp hydrogels with different crosslinker contents.

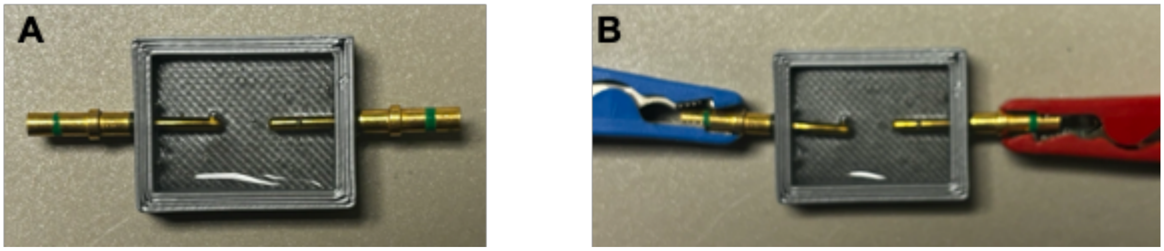

**Figure SI15.** (A) The impedance of GKAmp hydrogel was tested using a 3D printed mold and connected to a gold rod electrode. (B) The positive and negative electrodes were connected at both ends during the test.

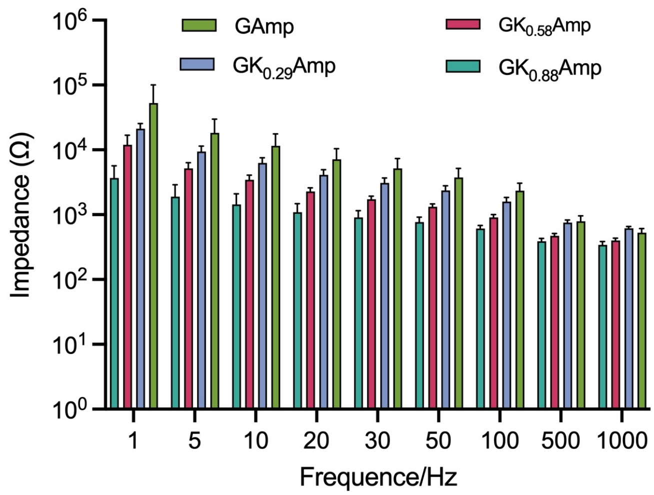


**Figure SI16.** Impedance values of GKAmp hydrogel tested at different KCl contents at corresponding frequencies

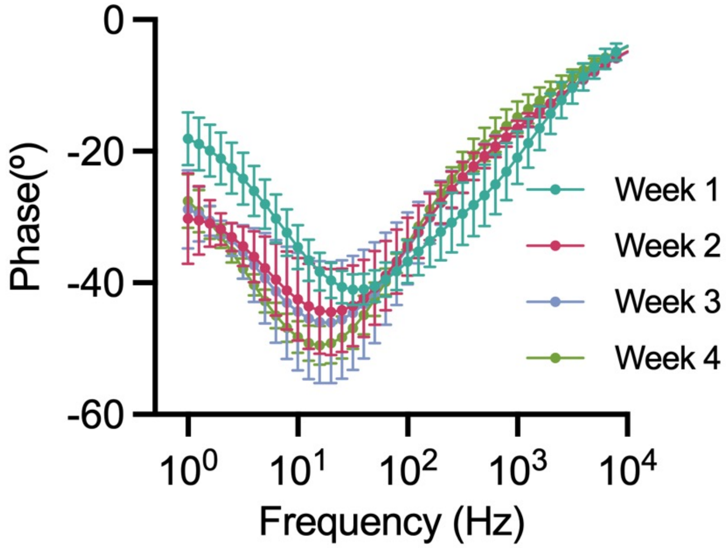


**Figure SI17.** Evolution of impedance frequency response and phase characteristics of GKAmp hydrogel over 1–4 weeks.

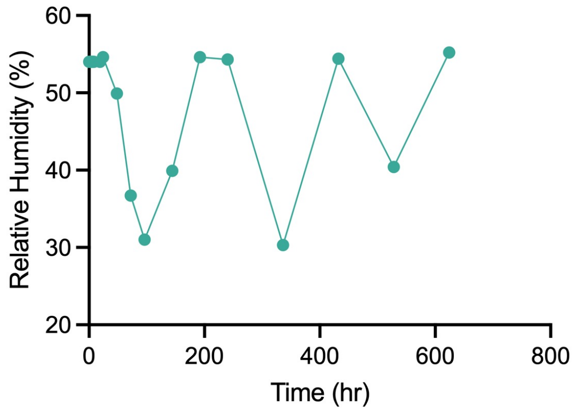


**Figure SI18.** Humidity variations during weight measurement over 0–600 hours.

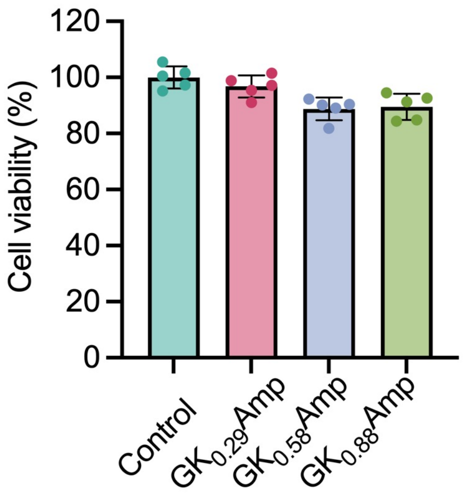


**Figure SI19.** Cell viability analysis for hydrogel with different KCl concentrations after 48 h of co-culture.

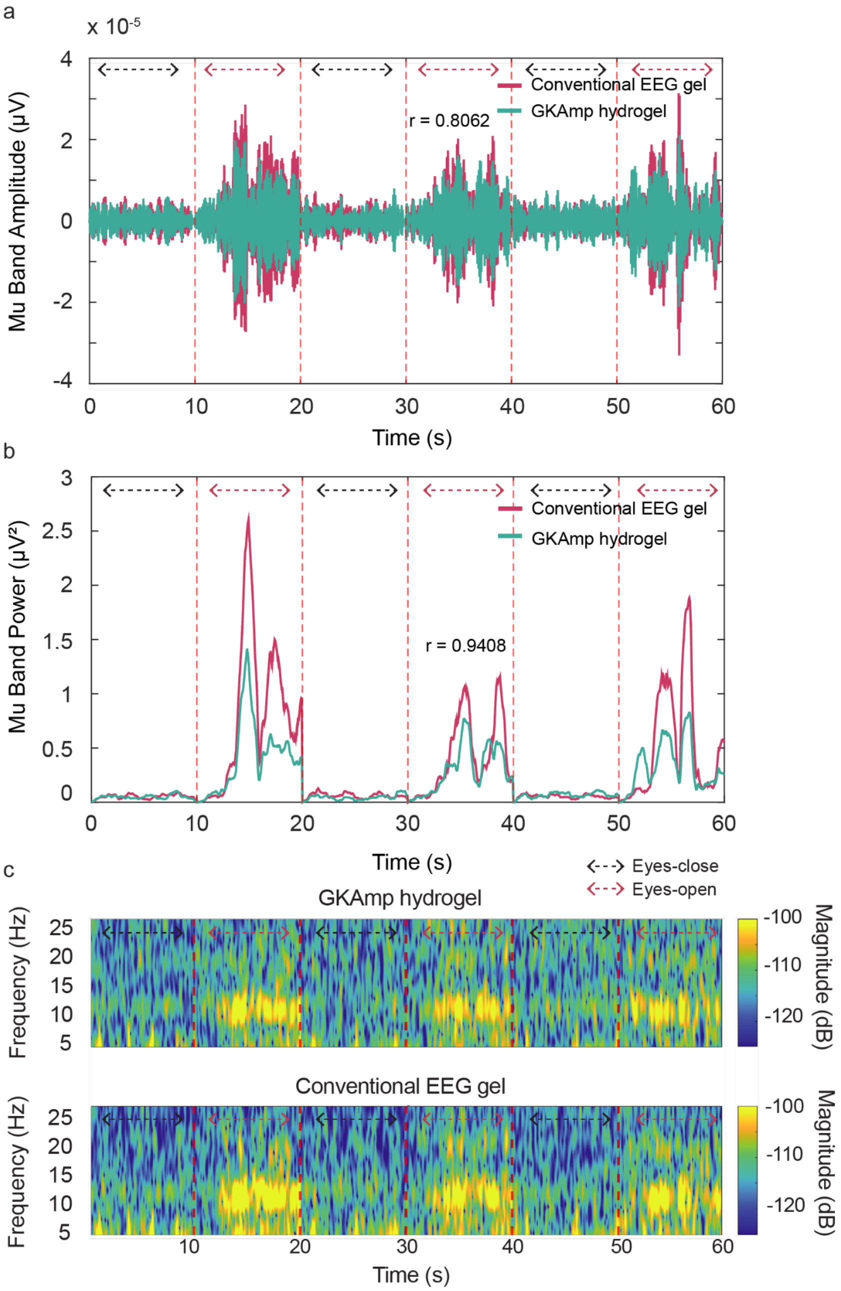


**Figure SI20. Comparison of EEG signal fidelity between GKAmp hydrogel and conventional EEG gel electrodes.**

(a) Mu-band amplitude signals recorded using GKAmp hydrogel (green) and conventional EEG gel (red) electrodes during alternating eyes-open and eyes-closed conditions. The high correlation coefficient (r=0.8062) demonstrates comparable signal quality between the two electrode types. (b) Mu-band power extracted from the same recordings showed a strong correlation (r=0.9408) between GKAmp and conventional gel electrodes, with consistent power peaks during eyes-closed periods. (c) Time-frequency spectrograms of EEG signals recorded using GKAmp hydrogel (top) and conventional EEG gel (bottom) electrodes, highlighting comparable spectral power distributions across eyes-open and eyes-closed conditions.

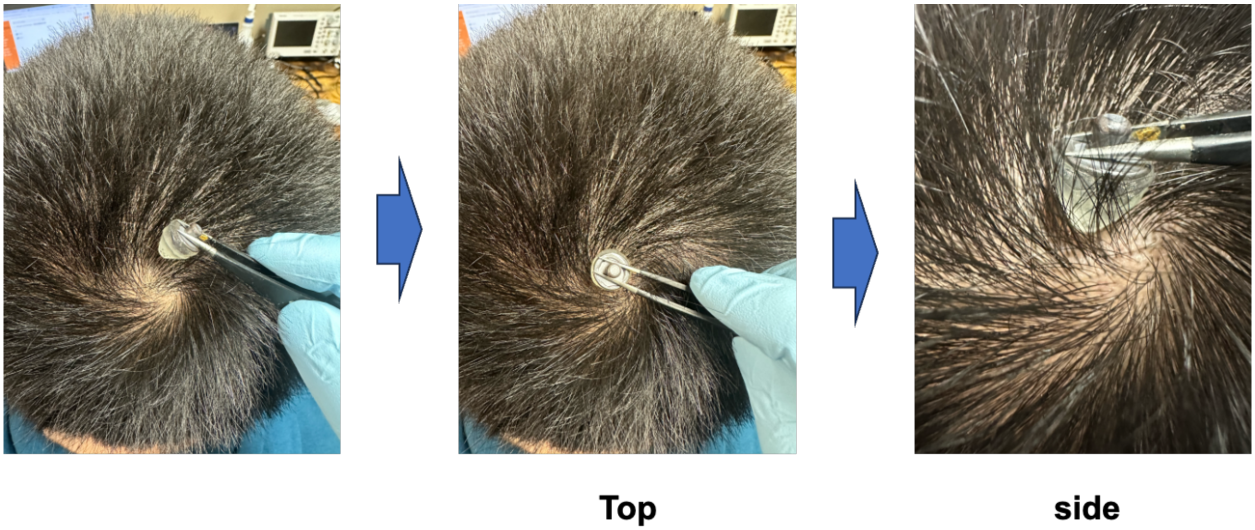


**Figure SI21.** Integration of hydrogel and 3D-printed elastic electrodes in contact with the scalp with the interference of hair. (a) Integration of hydrogel and 3D-printed porous conical elastic electrodes before and after contact with the scalp (b) top and (c) side photographs

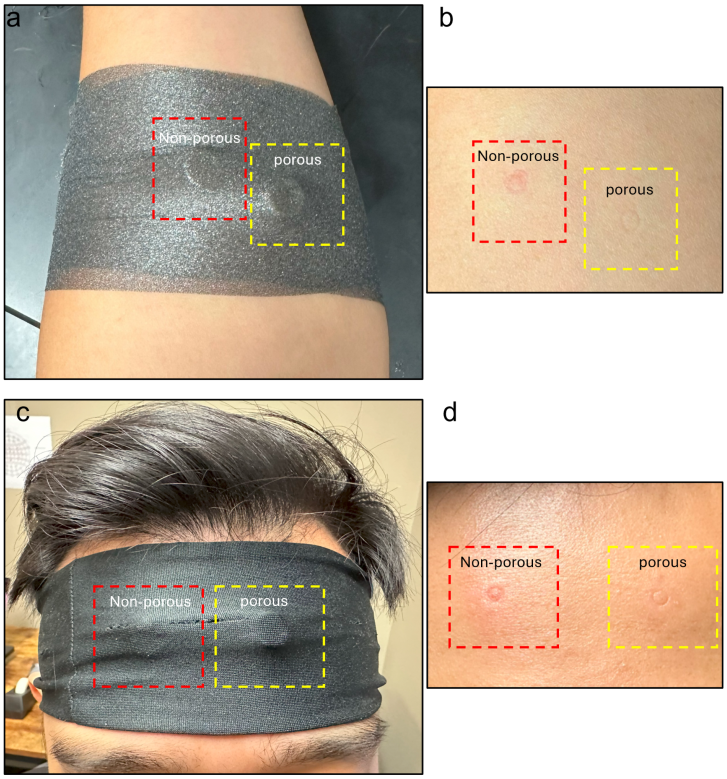


**Figure SI22. Comparison of residual redness caused by porous and non-porous platforms after 10 minutes of wear.**

(a) Testing on a human volunteer's arm with marked regions for non-porous (red dashed box) and porous (yellow dashed box) platforms. (b) Close-up view of residual redness on the arm after 10 minutes of wear, showing more pronounced redness for the non-porous platform compared to the porous platform. (c) Testing on a human volunteer's forehead with marked regions for non-porous (red dashed box) and porous (yellow dashed box) platforms. (d) Close-up view of residual redness on the forehead after 10 minutes of wear, again demonstrating more significant redness for the non-porous platform relative to the porous platform.

**Figures related to: MindStretcH Enables Long-Term and Reliable Motor Imagery EEG Monitoring**

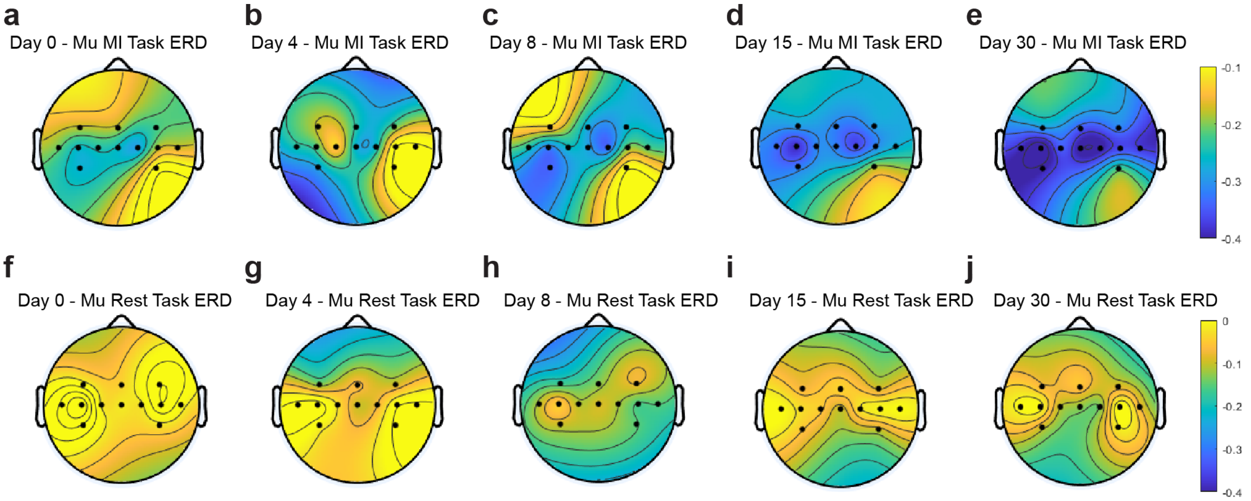


**Figure SI23. Topographical maps of mu-band (8-13 Hz) ERD during MI and rest tasks across 5 sessions.**

(a–e) Mu-band ERD during MI tasks over five sessions, showing progressive refinement of contralateral sensorimotor cortex activation, with stronger and more localized desynchronization in later sessions. (f–j) Mu-band ERD during rest tasks over five sessions, indicating minimal desynchronization and consistent baseline activity, highlighting the specificity of task-related modulation.

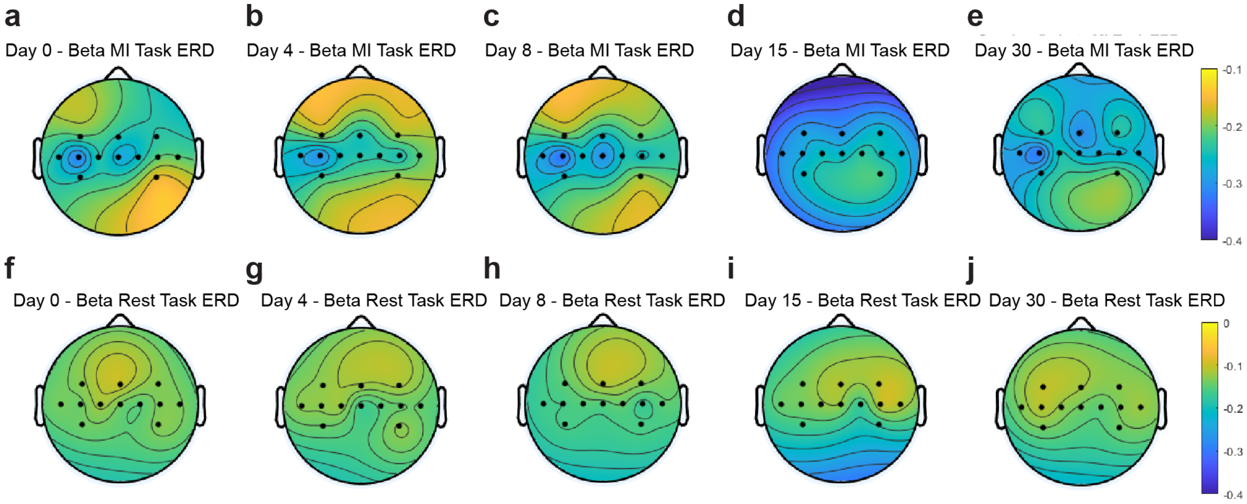


**Figure SI24. Topographical maps of beta-band (18-30 Hz) ERD during MI and rest tasks across 5 sessions.**

(a–e) Beta-band ERD during MI tasks over five sessions, showing broader and less localized desynchronization compared to the mu-band, with moderate task-specific modulation. (f–j) Beta-band ERD during rest tasks over five sessions, displaying diffuse and weak desynchronization, further emphasizing reduced task specificity in the beta-band compared to the mu-band.

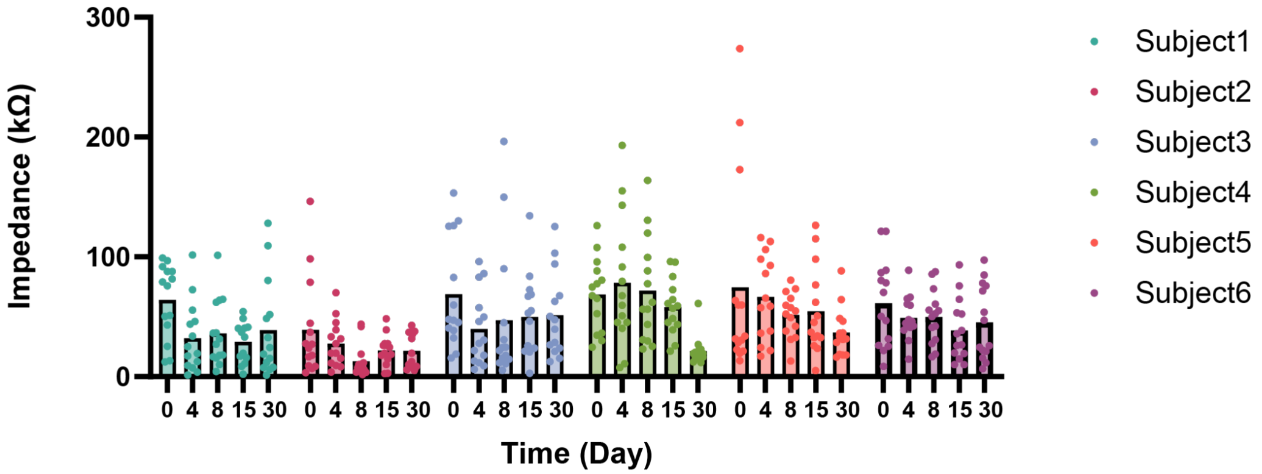


**Figure SI25. Subject-wise scalp-electrode impedance values across 5 sessions.**
Impedance values for all 6 subjects were measured at each session (days 1, 4, 8, 15, and 30).

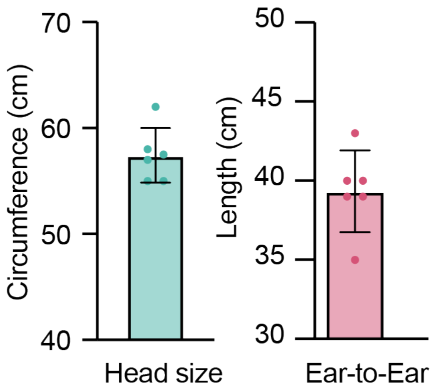


**Figure SI26. Anthropometric measurements of study participants.**

(a) Head circumference (mean = 57.4 cm) and (b) ear-to-ear length (mean = 39.3 cm) of the 6 participants (n = 6, female: male = 1:1). Unlike traditional EEG caps that failed to accommodate larger head sizes, MindStretcH successfully fit all participants, demonstrating its adaptability to diverse anthropometric characteristics.

**Figures related to: Machine Learning-Driven Multi-Week Closed-Loop Motor Imagery-based BCI Operation**

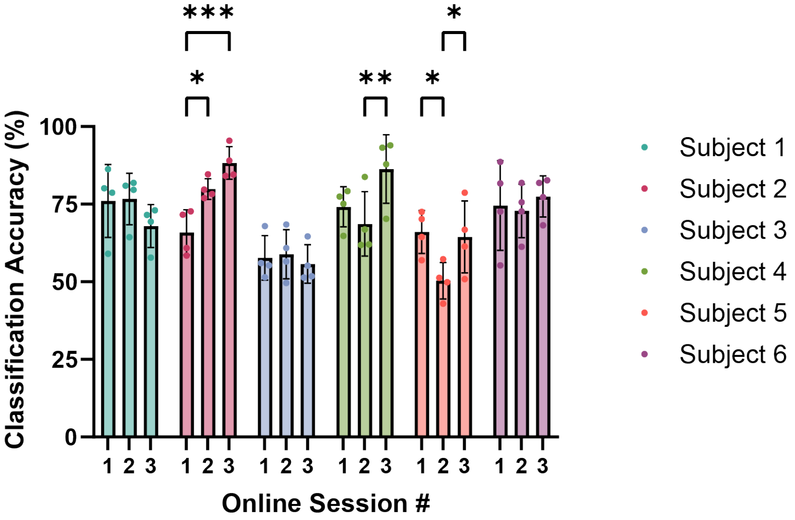


**Figure SI27. Subject-wise sample-wise classification accuracy across the three online sessions.** Bar graphs represent the subject-specific classification accuracy (%) for each subject (n=6) in sessions 1, 2, and 3. Error bars indicate standard deviation. Pairwise comparisons from the Tukey post-hoc test from two-way ANOVA are provided for significant effects observed (*p < 0.05, **p < 0.01, ***p < 0.001): Subject 2 (session 1 vs session 2, session 1 vs session 3), Subject 4 (session 2 vs session 3), and Subject 5 (session 1 vs session 2, session 2 vs session 3).

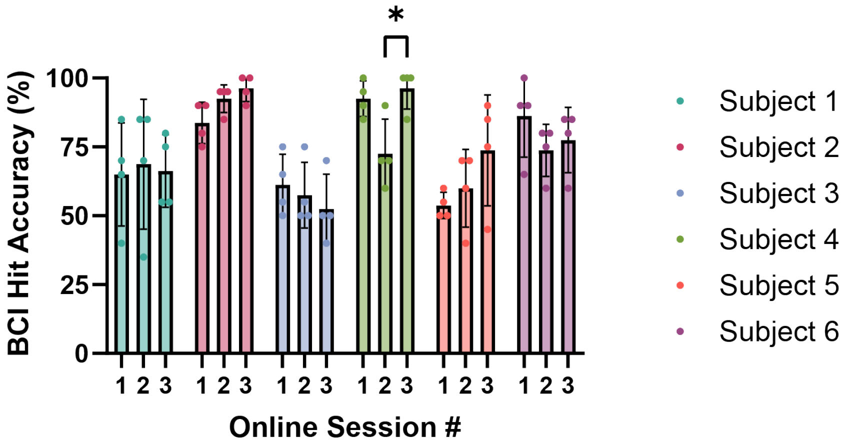


**Figure SI28. Subject-wise BCI hit accuracy across the three online sessions.**
Bar graphs represent the subject-specific BCI hit accuracy (%) for each subject (n=6) in sessions 1, 2, and 3. Error bars indicate standard deviation. Pairwise comparisons from Šídák's multiple comparisons post-hoc test from two-way ANOVA are provided for significant effects observed (*$p < 0.05$): Subject 4 (session 2 vs session 3).

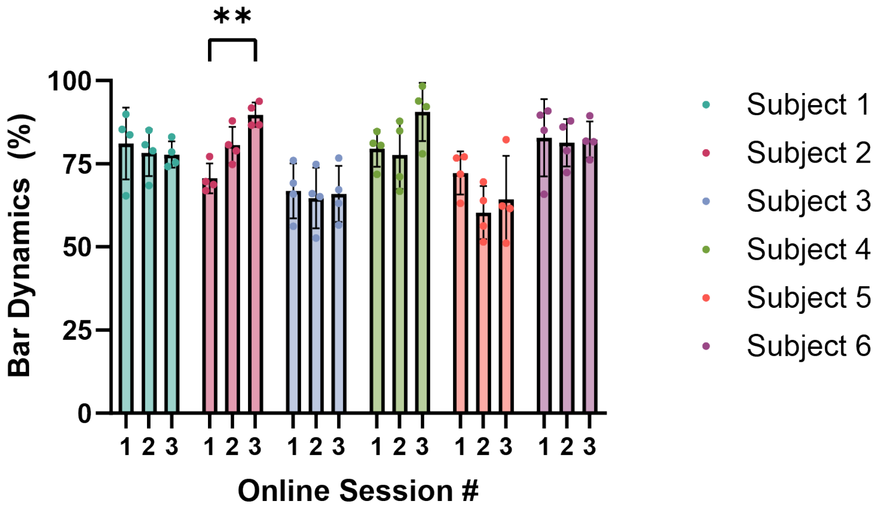


**Figure SI29. Subject-wise Bar Dynamics across the three online sessions.**
Bar graphs represent the subject-specific Bar Dynamics (%) for each subject (n=6) in sessions 1, 2, and 3. Error bars indicate standard deviation. Pairwise comparisons from Šídák's multiple comparisons post-hoc test from two-way ANOVA are provided for significant effects observed (**$p < 0.01$): Subject 2 (session 1 vs session 3).

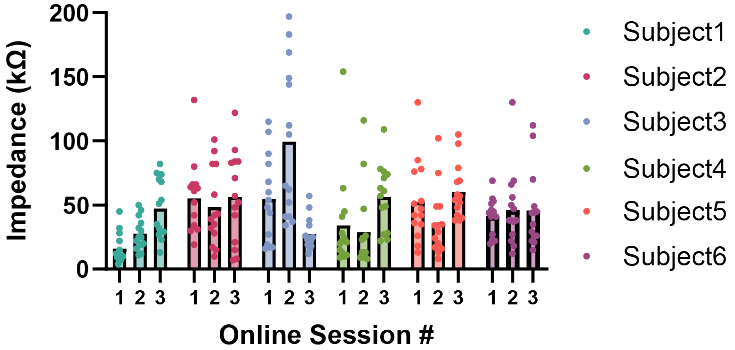


**Figure SI30. Subject-wise scalp-electrode impedance values across 3 online sessions.** Impedance values for all 6 subjects were measured at each session.
